# Structural Basis for OAS2 Regulation and its Antiviral Function

**DOI:** 10.1101/2025.01.28.635220

**Authors:** Veronika Merold, Indra Bekere, Stefanie Kretschmer, Adrian F. Schnell, Dorota Kmiec, Rinu Sivarajan, Katja Lammens, Rou Liu, Julia Mergner, Julia Teppert, Maximilian Hirschenberger, Sarah Hammes, Kathrin Buder, Marcus Weitz, Lars M. Koenig, Andreas Pichlmair, Nadine Schwierz, Konstantin M.J. Sparrer, Min Ae Lee-Kirsch, Carina C. de Oliveira Mann

## Abstract

Oligoadenylate synthetase (OAS) proteins are immune sensors for double-stranded RNA and critical for restricting viruses. OAS2 comprises two OAS domains, only one of which can synthesize 2’-5’-oligoadenylates for RNase L activation. Existing structures of OAS1 provide a model for enzyme activation, but do not explain how multiple OAS domains discriminate RNA length. Here, we discover that OAS2 exists in an autoinhibited state as a zinc-mediated dimer and present a mechanism for RNA length discrimination: the catalytically deficient domain acts as a molecular ruler that prevents autoreactivity to short RNAs. We demonstrate that dimerization and myristoylation localize OAS2 to Golgi membranes and that this is required for OAS2 activation and restriction of viruses that exploit the endomembrane system for replication, e.g. coronaviruses. Finally, our results highlight the non-redundant role of OAS proteins and emphasize the clinical relevance of OAS2 by identifying a patient with a loss-of-function mutation leading to autoimmune disease.

## INTRODUCTION

Double-stranded RNA (dsRNA) is one of the most relevant signatures of viral replication that is sensed by the immune system. Recognition of this pathogenic dsRNA in the cytoplasm occurs through different classes of immune receptors and restriction factors. 2′-5′ Oligoadenylate Synthetase (OAS) proteins are interferon-induced antiviral enzymes that require binding to dsRNA for activation ^1–4^. As part of the nucleotidyltransferase (NTase) protein family and structurally related to the dsDNA receptor cyclic GMP-AMP synthase (cGAS), OAS proteins synthesize unique second messengers that feature 2′-5′ phosphodiester bonds by oligomerizing ATP into 2′-5′-oligoadenylates (2’–5’OA) of varying lengths ^4–8^. 2’–5’OA in turn bind and activate downstream latent RNase L, which degrades both viral and cellular single-stranded RNA (ssRNA), resulting in translational arrest and ultimately restricting viral replication ^9–15^. A dysregulation of OAS1 function has been implicated in various autoinflammatory conditions and is associated with an increased risk of severe outcomes in COVID-19^16–23^. Recently, hypomorphic loss-of-function variants in both OAS1 and OAS2 have been connected to multisystem inflammatory syndrome in children (MIS-C) following SARS-CoV-2 infection ^24^.

Humans express three catalytically active OAS proteins, OAS1, OAS2 and OAS3. OAS1 consists of a single OAS domain of the polymerase beta (pol-β)-like NTase fold that harbors the catalytic triad required for 2’–5’OA synthesis ^25–28^. OAS2 and OAS3 are unusual members of the NTase protein family, as they contain two and three OAS domains, respectively, which have evolved through gene duplication events ^29–31^. Only one OAS domain in each OAS protein has retained the catalytic triad in the active site, raising the question of the role played by the catalytically inactive pseudo-OAS domains^32^. Despite not having catalytic activity these pseudo-OAS domains were shown to be required for activation of OAS2 as well as OAS3 ^33–36^. A current model suggests that OAS proteins have evolved to discriminate between different RNA lengths with OAS1 being activated by short >18 bp long dsRNA, OAS2 by >35 bp medium long and OAS3 by >50 bp long dsRNA^37^. While the structural mechanism for OAS1 activation by short dsRNA is well described, a major limitation in our understanding of OAS protein function is the lack of structural information for multi-domain OAS proteins. All current structural and mechanistic information is therefore derived from crystal structures of isolated OAS domains such as OAS1 or OAS3 domain I (DI) ^26–28,33^. Thus, despite being among the first antiviral proteins discovered, the mechanism underlying the length-dependent activation of OAS proteins by dsRNA remains elusive. Importantly, in addition to different RNA ligands, the varied subcellular localization of OAS proteins has recently been demonstrated to shape their antiviral function and contribute to their unique roles in virus restriction. While OAS3 recognizes viruses that replicate in the cytosol, OAS1 isoform p46 is targeted to membranes of the endomembrane system through prenylation and has evolved to detect viral dsRNA directly at their replication organelles ^19,20,38–41^. Similar to OAS1, OAS2 has been shown to be myristoylated at its N-terminus ^42^. However, the impact of subcellular targeting of OAS2 on its specificity and antiviral activity remains unclear. These unexplored questions regarding OAS protein function have resulted in a generalized understanding of their roles, often viewed solely through the lens of RNase L activation. However, we are now beginning to appreciate the non-redundant functions of OAS proteins in antiviral immunity. Here we define an unexpected mechanism of OAS2 regulation that prevents OAS2 activation by short dsRNA and determine how subcellular localization of OAS2 defines its antiviral activity. We solved the dimeric structure of OAS2 in an autoinhibited state where the catalytically active OAS2 domain DII is trapped in an inactive conformation. Dimerization of OAS2 allows the catalytically deficient domain DI to measure RNA length explaining how OAS2 requires longer RNAs for activation than OAS1. OAS2 antiviral activity is dependent on association of OAS2 to Golgi membranes via myristoylation. In line with non-redundant functions of OAS proteins, complete loss of function of OAS2 leads to severe autoimmune disease in a patient. Taken together, our results define a mechanism of OAS2 regulation that explains how OAS2 discriminates between RNA lengths and how association with Golgi membranes is required for OAS2 antiviral functions.

## RESULTS

### Structure of human OAS2 reveals dimerization via a zinc binding site

OAS2 consists of two OAS domains: a catalytically deficient domain DI (residues 13-325) and a catalytically active domain DII (residues 345-686), which share their overall structural fold with limited sequence identity (42.9%). Both domains are predicted to be connected by a flexible linker (residues 325-345) and flanked by a flexible N-(10 aa) or C-terminus (36 aa). (Fig.1A, SI Fig.2.). We purified full-length OAS2 p71 to understand how multiple domains connected by a linker are regulated.

**Figure 1.**
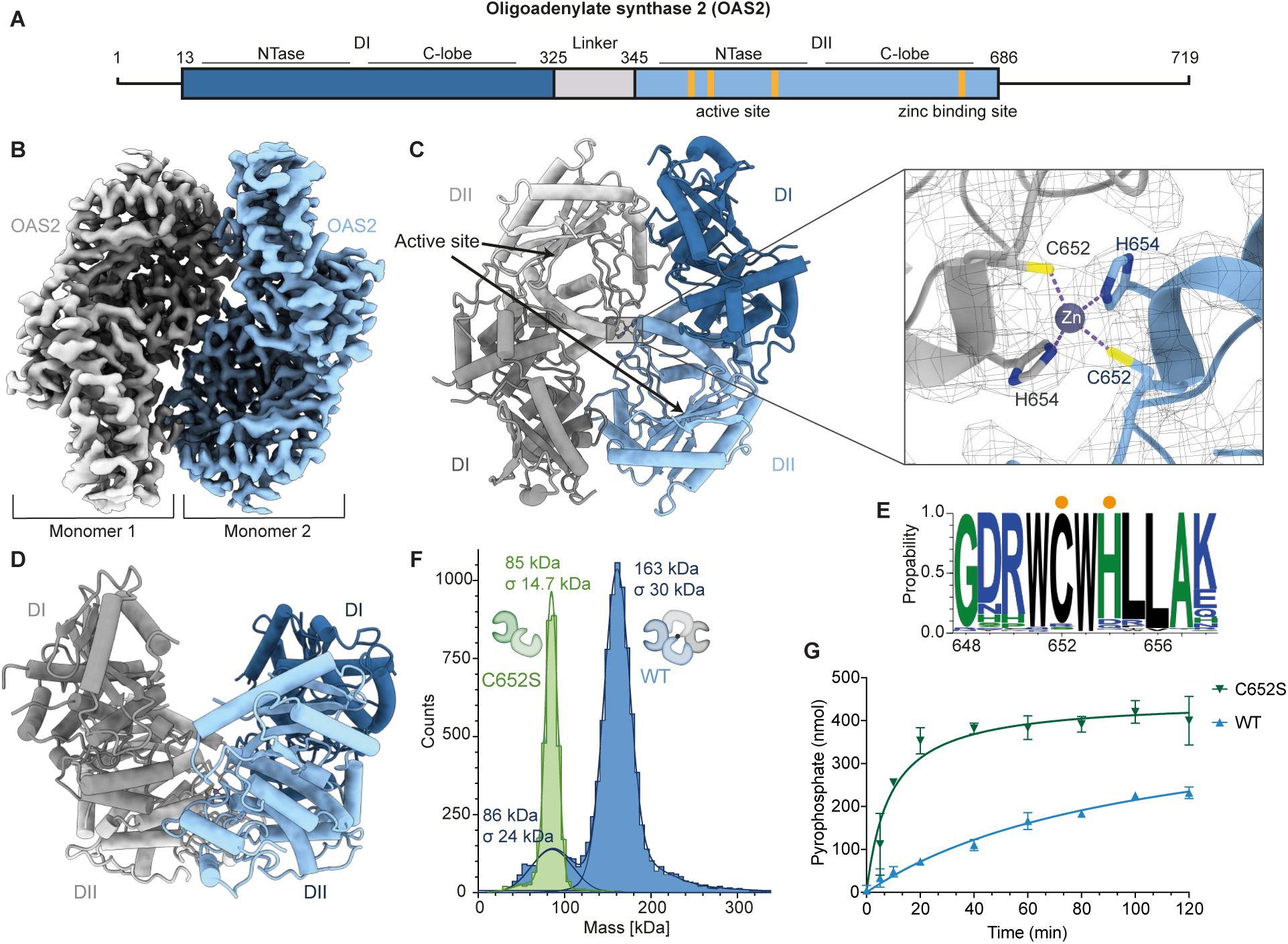
Structure of human OAS2 reveals dimerization via a zinc binding site. (A) Domain organization of human OAS2. (B) Cryo-em reconstruction of OAS2 in 3.3 Å resolution. Two monomers are depicted in blue and grey, respectively. (C) Atomic model of OAS2 dimer with close-up of zinc binding site. Monomers are depicted in blue and grey, respectively. Inactive domain DI is depicted in darker color, active domain DII is depicted in lighter color. Close-up view shows Zn^2+^ coordination of DII C652 and H654 from each monomer. Electron density is depicted as grey mesh. (D) Side view of OAS2 dimer. (E) Sequence logo of 210 species showing conservation of C652 and H654. (F) Mass photometer analysis shows that OAS2 C652S is monomeric (green) compared to dimeric OAS2 wild-type (blue). (G) *In vitro* chromogenic activity assay of 100 nM OAS2 wild-type (blue) and C652S (green) with 82bp long dsRNA (100 nM) (mean ± SD of n = 3). Monomeric OAS2 C652S shows increased activity compared to dimeric OAS2.

**Figure 2.**
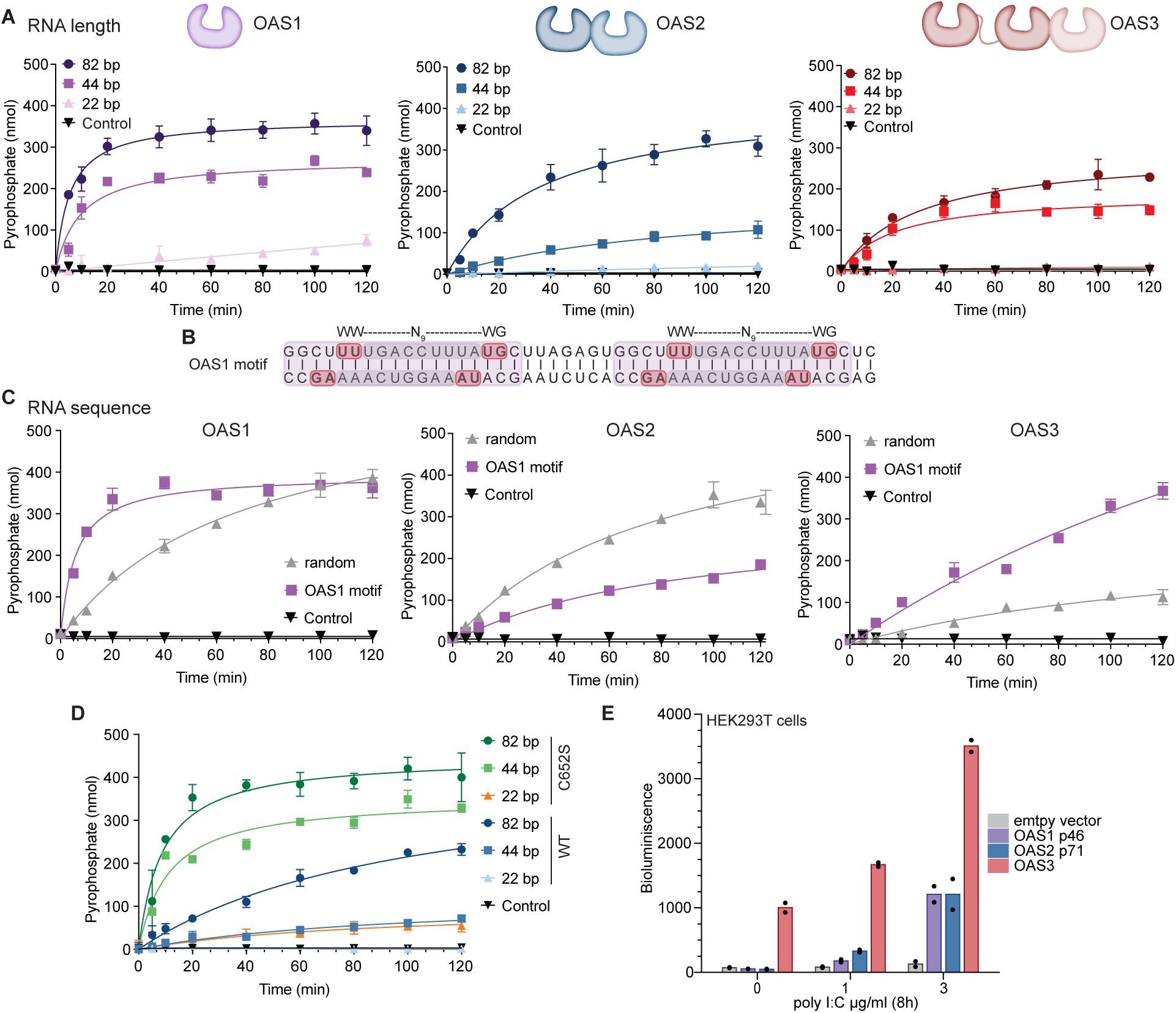
RNA ligand requirements for the activation of OAS2 differ from OAS1 and OAS3. (A) *In vitro* chromogenic activity assay of 200 nM OAS1 p46, OAS2 p71 and OAS3 p100 each with 22 bp, 44 bp, and 82 bp dsRNA (200 nM) (mean ± SD of n = 3). Comparison of OAS protein activation by different RNA lengths. (B) Sequence of preferred OAS1 motif. WWN9WG motif is colored in red. Repetitive sequence is colored in purple. (C) Sequence preference of OAS proteins. *In vitro* assay as in (A) with of 200 nM OAS1, OAS2 and OAS3 with 44 bp dsRNA (200 nM) containing OAS1 motif or random sequence (mean ± SD of n = 3). RNA containing the OAS1 motif (purple) activates strongly OAS1 and OAS3. In contrast, OAS2 shows a preference for the RNA lacking this motif (random). (D) *In vitro* assay as in (A) with 100 nM OAS2 wild type (blue) and OAS2 C652S (green, orange) with 23 bp, 44 bp, and 82 bp dsRNA (100 nM) (mean ± SD of n = 3). OAS2 C652S can be activated by short dsRNA in contrast to OAS2 wild-type. (E) Comparison of OAS protein activity in cells with 2’-5’OA biosensor for OAS1, OAS2 and OAS3. OAS3 revealed highest levels of 2’-5’OA synthesis and was active even in the absence of poly I:C treatment. HEK293T RNASEL KO cells were transiently transfected with different OAS proteins for 48h followed by treatment with poly I:C. Bioluminescence was measured at 8h post poly I:C treatment. Assay is representative of at least three independent experiments. Bars represent means of technical replicates (dots).

During the purification of the OAS2 protein, gel filtration showed that OAS2 is a dimer, in contrast to monomeric OAS1 and OAS3 (SI Fig.1A). Oligomerization was already found to be important for OAS1 and OAS2 ^34,36,43^. To mechanistically understand how dimerization of OAS2 regulates its activation, we solved the structure of human OAS2 using cryo-electron microscopy (cryo-em). 2D classification and 3D reconstruction revealed an OAS2 dimer at 6.2 Å (SI Fig. 1B, C). Subsequent homogenous and non-uniform refinements further improved overall resolution of 3.3 Å (SI Fig. 1C-F). The OAS2 dimer has a bowl-shaped conformation with an interface of 3441.9 Å^2^ between the monomers, which are positioned in an antiparallel orientation to each other (Fig. 1B, C, D). This mode of dimerization traps OAS2 in an inactive conformation, with the active site facing the inside of the dimer interface (SI Fig. 3D, Fig.1C). In this conformation, the active site is not aligned, but the helical P-loop (residues 396-399) is formed in the absence of dsRNA, as has been reported for other NTase/ OAS domain structures (SI Fig. 3D)^27,33,44,45^. This places DII within the OAS2 dimer in a hybrid conformation. Our structure also revealed that the linker (residues 326-345) connecting the two OAS domains is surprisingly well resolved (SI Fig. 3A), as are the C-terminal residues (670-684) flanking OAS2 DII, suggesting that these residues keep the two OAS domains in close proximity, limiting their flexibility towards each other.

**Figure 3.**
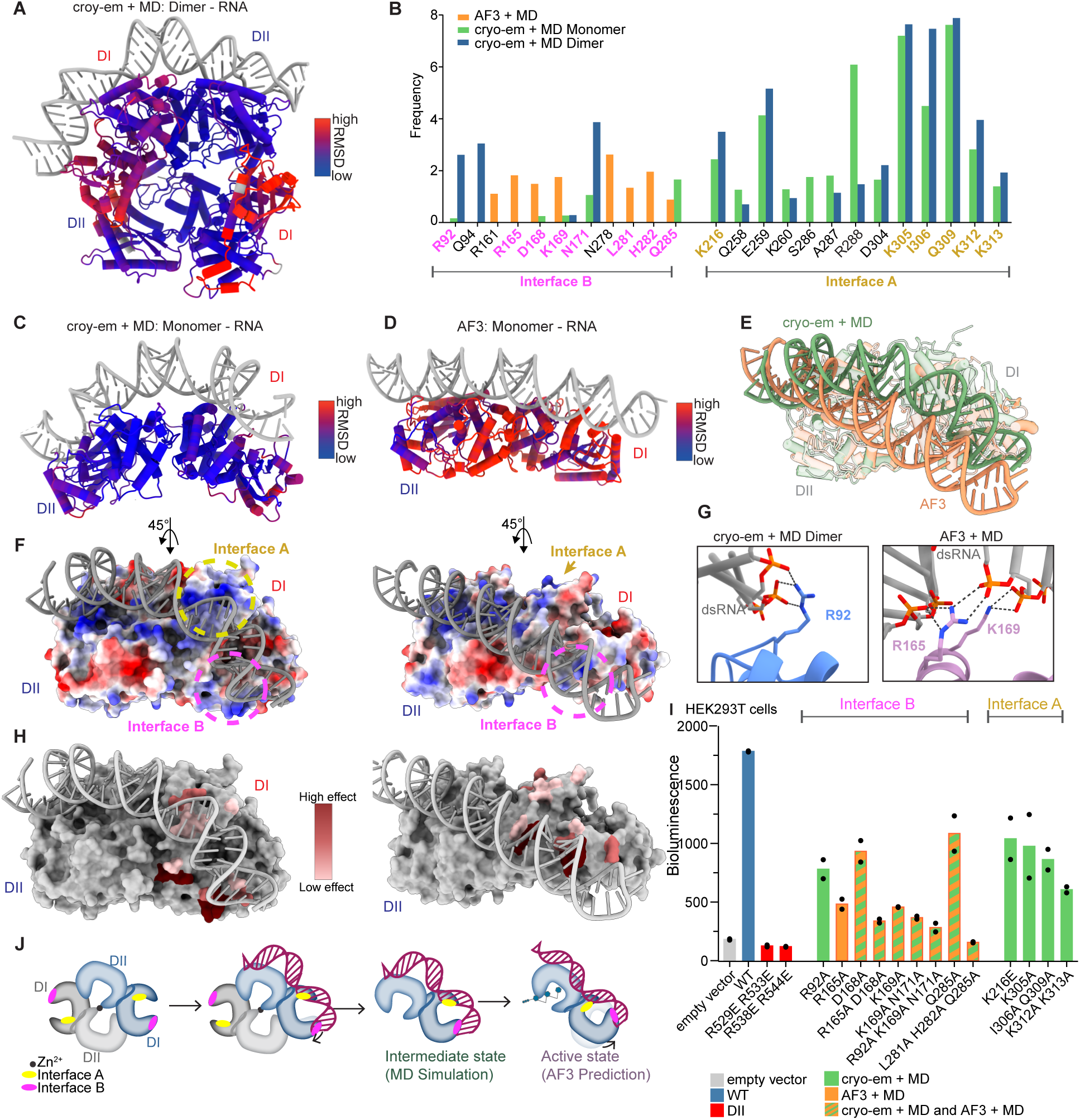
OAS2 DI functions as a regulatory domain that measures RNA length using a non-canonical interface. (A) MD simulation of OAS2 dimer in complex with dsRNA (grey), colored by RMSD values using the cryo-em structure as a reference. High RMSD values are represented in red, while low values are shown in blue. (B) Contact frequencies of DI with dsRNA, calculated based on cryo-em+MD and AF3+MD approaches. Residues tested in panel (I) are highlighted in bold and color-coded. (C) MD simulation of OAS2 monomer in complex with dsRNA (grey). Structure is colored as described in (A). (D) AF3 prediction of OAS2 monomer in complex with dsRNA (grey). Structure is colored as described in (A) (E) Superposition of OAS2 with dsRNA from cryo-em+MD and AF3+MD approaches in green and orange, respectively. (F) Electrostatic surface representation from cryo-em+MD of monomeric OAS2 in complex with dsRNA (left) and AF3 prediction (right). Interfaces A and B are highlighted with yellow and pink dashed circles, respectively. (G) Close ups of Interface B of the cryo-em+MD (left) and AF3 predicted structure (right). (H) Surface view of structure depicted in (F), with color code indicating the effect of mutations as shown in (I). (I) Analysis of OAS protein activity in cells with 2′-5′OA biosensor for OAS2 RNA-binding site mutants showing reduced activity for OAS2 mutants in non-canonical RNA-binding sites in DI. HEK293T cells were transiently transfected with different OAS2 constructs for 48h followed by treatment with poly I:C for 24h. Assay is representative of three independent experiments. Bars represent means of technical replicates (dots). (J) Schematic overview of the RNA binding mechanism of OAS2. The OAS2 dimer is auto-inhibited, and upon RNA binding, it monomerizes. The MD simulation illustrates an intermediate state, while the AF3 prediction more accurately represents the active state bound to RNA. RNA binding interfaces A and B are colored in yellow and pink respectively.

The OAS2 dimer is held together by an unexpected tetrahedral zinc coordination site in the middle of the hydrophobic dimerization interface (Fig. 1C). Residues C652 and H654 of the catalytically active DII of each monomer coordinate the binding of a zinc atom. This DII-driven dimerization mechanism allows DII to remain inactive even in the absence of DI, explaining why previous attempts to activate the isolated DII domain *in vitro*, analogous to OAS1, have failed ^34,36^. C652 and H654 are conserved in OAS2 but not in OAS1 and OAS3, revealing a dimerization mechanism unique to OAS2 (Fig.1E, SI Fig.2). When comparing the Alphafold3 ^46^ prediction for the OAS2 dimer with the model of our structure, we observe large deviations (RMSD = 1.01 nm). Even when provided with information about Zn^2+^ as a ligand, the resulting AF3 prediction did not evidence the same conformation as our OAS2 structure (RMSD = 0.68 nm) (SI Fig.3B, C).

Mutation of OAS2 C652S is sufficient to disrupt the dimer interface and to render OAS2 monomeric (Fig. 1F, SI Fig.1A). Given the inactive conformation of the active site of the OAS2 dimer, we hypothesised that OAS2 must monomerize to be properly activated by its ligand dsRNA. To test this hypothesis, we compared the activity of the wild-type OAS2 with that of the OAS2 C652S using an established *in vitro* colorimetric assay that monitors the release of pyrophosphate, a by-product of OAS2 enzymatic activity. Our results show that the OAS2 C652S is significantly more active in the presence of dsRNA than the wild-type OAS2, providing evidence that monomerization of OAS2 is a previously unknown requirement that facilitates activation by dsRNA. (Fig. 1G). These results provide the structural framework for multiple OAS domains and reveal a mechanism by which dimerization via metal coordination keeps the protein in an auto-inhibited state, creating an unexpected level of regulation that controls 2’-5’ oligoadenylate synthesis.

### RNA ligand requirements for the activation of OAS2 differ from OAS1 and OAS3

Research into how OAS proteins recognize RNA has concentrated on their ability to discriminate dsRNAs based on length^47^. OAS1 binds dsRNA to a positively charged groove opposite the active site with a footprint of 17 bp and with a preference for a consensus sequence that allows it to contact dsRNA at two segments of the minor groove of the dsRNA helix ^27,28,37^. Lack of structural information for multi-domain OAS proteins, lead to the current model in which the mode of RNA recognition is extrapolated from OAS1 to other members of the OAS family. Longer RNA must bridge and bind to the same positively charged groove on multiple OAS domains to activate the respective enzyme. To understand if and how the ligand requirements of OAS2 differ from those of other OAS proteins, we first analysed the effects of RNA length dependence for the activation of OAS1, OAS2 and OAS3 (Fig.2A). In agreement with previous experiments, OAS1 is active when incubated with 22 bp long dsRNA and its activity increases gradually with longer RNAs ^27,28,37,48^. In contrast, OAS2 is inactive with 22 bp long dsRNA and shows minimal activity with 44 bp. Interestingly, the OAS2 enzyme exhibits proper activation only with long dsRNA of 82 bp. OAS3 on the other hand is active in the presence of RNA with 44 bp length and its activity does only slightly increase even when incubated with longer RNAs. Isoforms OAS1 p42 and OAS2 p69 demonstrated comparable activation behaviour, although with slightly reduced activity levels (SI Fig.4A). Accordingly, all subsequent experiments were conducted with the OAS2 isoform p71. Our results demonstrate that *in vitro* OAS2 requires longer RNAs than OAS3 for proper activation, supporting the hypothesis that the activation mechanism is more complex than the current model suggests. To understand how OAS2 dimerization regulates activation by RNA, we next repeated this activity assay with OAS2 C652S (Fig. 2D). As shown in Fig.1G, OAS2 C652S is significantly more active than OAS2 in the presence of long dsRNA. Notably, OAS2 C652S is even activated by dsRNA of 22 bp length in contrast to the wild-type protein, suggesting that binding of RNA to the catalytically active OAS2 DII is sufficient for activation. These results demonstrate that OAS2 DII has a comparable activation mechanism to OAS1 in the presence of short RNAs >20 bp, when not trapped in the inactive dimeric conformation. A conserved RNA motif WWN_9_WG (where W can be either A or U, and N can be any nucleotide) was demonstrated to strongly activate OAS1 (Fig.2B) ^27,28,49–52^. To determine whether OAS2 or OAS3 have the same sequence requirements as OAS1, we performed activity assays with two different RNA sequences. We compared a 40 bp long dsRNA with the duplicated conserved WWN_9_WG motif (OAS1 motif) and one dsRNA containing a random sequence (SI Fig. 4B). As expected, OAS1 activation was increased when incubated with 40bp RNA containing the WWN_9_WG motif (Fig.2C). Additionally, OAS3 showed similar behaviour to OAS1, with the OAS1 motif activating the protein better than the random sequence. Interestingly, activation of OAS2 was stronger with random RNA sequence than with the OAS1 motif, confirming that OAS2 has different requirements for its ligand RNA sequence.

**Figure 4.**
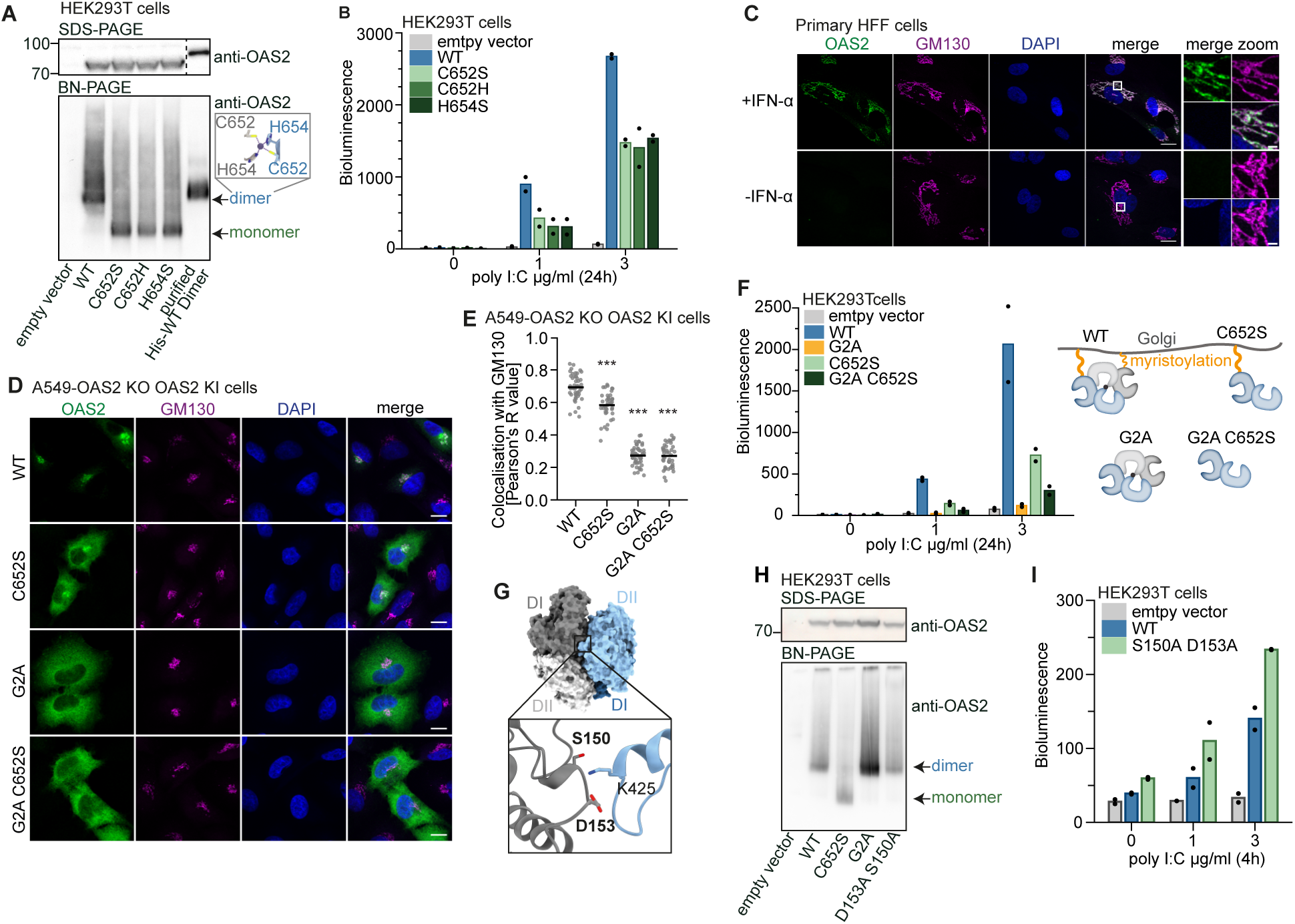
OAS2 dimerization and localization to the Golgi membrane via myristoylation is required for activation. (A) SDS-PAGE (top) and BN-PAGE (bottom) of different OAS2 constructs transiently expressed in HEK293T cells. Purified His-tagged OAS2 WT was loaded as a control for OAS2 dimerization. Zinc coordination is depicted on the right. (B) Analysis of OAS2 wild-type and monomeric mutants activity in cells with 2′-5′OA biosensor showing higher activity of dimeric wild-type than the monomeric mutants. HEK293T were transiently transfected with different OAS2 constructs for 48h followed by transfection of poly I:C. Bioluminescence was measured at 24h after poly I:C transfection. (C) Immunofluorescence Airyscan microscopy of endogenous OAS2 localization in primary HFF cells stimulated with 1000 U/ml with IFN-α for 24h. Cells were stained for OAS2 (green), golgi marker GM130 (magenta) and DAPI (blue). Scale bars represent 20 µm and 2 µm for merge zoom. (D) As in (C) for A549 OAS2 KO cells reconstituted with doxycycline-inducible OAS2 constructs treated with 1 µg/ml doxycycline for 24h. Scale bars represent 10µm. (E) Quantification of colocalization of OAS2 constructs and golgi marker GM130 from (D) based on Pearson correlation. Lines represent means from measurements of individual cells (dots). Statistical analysis was performed with Student’s T-test with Welch’s correction. *p < 0.05, **p < 0.01, ***p < 0.001. (F) Analysis of OAS2 activity in cells as in (B) for OAS2 wild-type and mutants with different localization showing that Golgi targeting is essential for the activity. Schematics depicting oligomeric state and localization of different OAS2 mutants are shown on the right. (G) Surface view of OAS2 dimer with each monomer depicted in grey and blue. Close up shows dimer interface interactions of S150 and D153. (H) SDS-PAGE and BN-PAGE analysis as in (A). (I) Analysis of OAS2 activity in cells as in (B) for OAS2 wild-type and DI-DII interaction mutant S150A D153A. Dimeric OAS2 wild-type conformation is destabilized in S150A D153A mutant increasing OAS2 activity. In (B), (F) and (I) assays are representative of three independent experiments. Bars represent means of technical replicates (dots).

To investigate OAS protein activation in cells, we used a previously established biosensor for 2’-5’OA ^53^. At first, it is evident that OAS3 produces more 2’-5’OA than OAS1 or OAS2, in line with previous reports ^39,40,54–56^ (Fig.2E, SI Fig.4C, D). Importantly, overexpression of OAS3 led to high 2’-5’OA synthesis by OAS3 in the absence of additional RNA stimulation. In contrast, OAS1 and OAS2 only became active after the cells had been treated with poly I:C. This suggests that OAS3 may be activated by endogenous dsRNA sources in HEK293T or is not in an auto-repressed state. The isoforms OAS1 p42 and OAS2 p69 produced less 2’-5’OA, which is consistent with our *in vitro* experiments (SI Fig.4A, C). Taken together, these results indicate that the current activation model does not explain the differences observed for OAS2 RNA-mediated activation.

### OAS2 DI functions as a regulatory domain that measures RNA length using a non-canonical interface

Superposition of the OAS1–RNA complex structure onto DI and DII of the OAS2 dimer reveals that proposing an RNA binding site spanning both OAS2 domains via the canonical binding interface is not feasible for two reasons (SI Fig.5B). First, the distance between the two interfaces cannot be extended by any 40 bp long dsRNA. Second, the most notable feature of this interface on DI is the presence of a negatively charged surface, which is unfavourable for the binding of RNA (SI Fig. 5A). This indicates that OAS2 DI binds dsRNA either through a different interface or, alternatively, does not bind RNA at all. To investigate how OAS2 DII and DI each contribute to the activation of OAS2 by RNA, we purified a construct of OAS2 DI. In agreement with previous reports, our experiments showed reduced binding to 82 bp long RNA for OAS2 DI, in contrast to significant binding of full-length OAS2 to dsRNA (SI Fig. 5C). These results prompted us to further investigate into the unknown contributions to RNA binding and functional role of the DI in the activation mechanism of OAS2.

**Figure 5.**
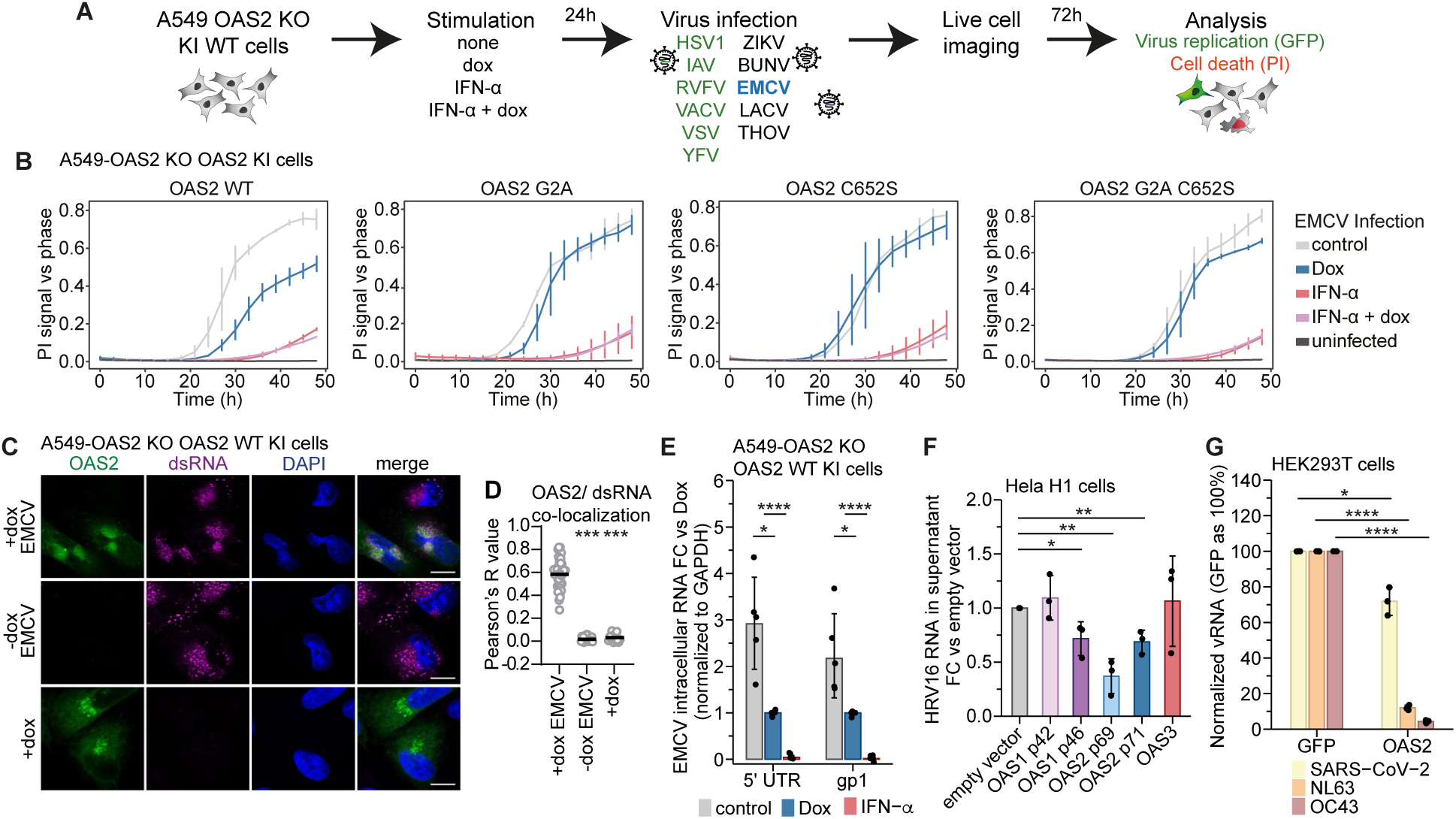
OAS2 restricts viruses replicating at the endomembrane system. (A) Experimental setup for virus screen in A549 OAS2 KO cells reconstituted with doxycycline inducible OAS2 WT. Cells were treated with 1 µg/ml doxycycline, 10U/ml IFN-α or combination of both for 24h. Afterwards cells were infected with a panel of GFP-reporter viruses (green) or non-reporter virus (black) described in Methods. GFP intensity for virus replication (GFP-reporter virus) and cell death with propidium iodide staining were measured using live cell imaging system Incucyte over time course of 72h. (B) Cell death in EMCV-infected (MOI 0.3) A549 OAS2 KO cells reconstituted with doxycycline-inducible OAS2 constructs. Cells were treated as described in (A). Cell death was quantified as area of propidium iodide signal versus cell confluency. Assays are representative of three independent experiments. Data are plotted with error bars representing the SD of the mean from two technical replicates. (C) Immunofluorescence Airyscan microscopy of OAS2 WT and viral dsRNA during EMCV infection. A549 OAS2 KO cells reconstituted with doxycycline-inducible OAS2 WT were treated with 1 µg/ml doxycycline for 24h followed by infected with EMCV for 24h with MOI 0.3. Cells were stained for OAS2 (green), dsRNA (magenta) and DAPI (blue). Scale bars represent 10 µm. (D) Quantification of OAS2 and dsRNA colocalization from (C). Lines represent means from measurements of individual cells (dots). Statistical analysis was performed with Student’s T-test with Welch’s correction. ***p < 0.001. (E) RT-qPCR analysis of intracellular EMCV RNA levels for 5’ UTR and gp1 in A549 OAS2 KO cells reconstituted with doxycycline-inducible OAS2 WT. Cells were treated with 1 µg/ml doxycycline or 10U/ml IFN-α for 24h followed by EMCV infection for 24h with MOI 0.3. (F) RT-qPCR analysis of HRV16 viral RNA levels in Hela H1 cell supernatants. Cells were transiently transfected with different OAS constructs and infected with HRV16 for 5 days. (G) RT-qPCR analysis of coronavirus RNA levels in culture supernatants after 72h of infection in HEK293T cells overexpressing OAS2 WT. In (D), (E) and (F) bars represent means of at least three independent experiments (dots), error bars show standard deviation. Statistical significance was calculated using paired T-test in panels (D), (F) or unpaired T-test in panel (E). * p<0.05, ** p<0.001, ***p < 0.001, **** p<0.0001.

To characterize the conformational dynamics of OAS2 and to predict the RNA binding site, we performed all-atom molecular dynamics (MD) simulations. The OAS2 dimer obtained from high-resolution cryo-em remains stable and close to the experimental structure (RMSD = 0.40 ± 0.06 nm) and shows low flexibility (SI Fig. 5D). The monomeric OAS2 protein exhibits much higher flexibility (SI Fig. 5D). Next, we used all-atom MD to predict the key residues involved in binding of dsRNA to OAS2. For the starting structure, the RNA was aligned to the binding site on DII of the cryo-em OAS2 structure, which was deducted from the alignment with the OAS1-RNA complex ^27,28^. We performed conformational sampling at high and physiological temperature and identified stable OAS2–RNA complexes. The results suggest that the RNA strongly bends along the dimer to span across DII and DI and introduces conformational changes at the interface connecting both monomers (Fig. 3A). The identified key residues from this cryo-em+MD approach relevant for the interactions of DI are depicted in (Fig. 3B). While the residues identified on DII are in line with the OAS1–RNA binding site, binding of dsRNA on DI occurs through a completely distinct interface, uncovering a non-canonical binding site specific to DI (Fig. 3A, B, SI Fig. 5A, E, F). Given that the RNA binding site on DII is expected, we focused on the interactions between DI and RNA in the subsequent analysis. Considering our previous findings that OAS2 is more active as a monomer, we performed further MD simulations focusing on the OAS2 monomer (Fig. 3C, SI Fig. 5D). As a complementary approach, we used AF3 to predict the structure of the OAS2-RNA complex (Fig. 3D, E) ^46^. The AF3 predictions for the monomeric OAS2 showed good agreement for the individual domains I (RMSD = 0.26 nm) and II (RMSD = 0.45 nm) but the overall structure deviated by 0.72 nm, indicating a conformational change in the orientation of the two domains. Therefore, we combined further MD simulations with the AF3 predictions (AF3+MD approach), to identify additional key residues involved in the binding of RNA to OAS2 DI. To better illustrate the differences, we classified key residues of DI with higher contact frequencies to RNA into two distinct interfaces, labeled A and B (Fig. 3B, F, G, SI Fig. 5G). RNA binding interface A, situated in closer proximity to DII, is constituted by residues present within the C-terminal α-helix bundle (DI-α14, DI-α15 and the loop between DI-14 and DI-α15). RNA binding interface B is comprised of residues of the N-terminal lobe situated DI-α5 and on loops between DI-α9 and DI-α10. The AF3+MD approach resulted in a less bent RNA compared to the results from the cryo-em+MD method and showed more contacts with residues on DII (Fig. 3C, D). While the former does not indicate RNA contacts with residues in interface A, it does reveal interactions between the RNA and residues in the DI RNA-binding interface B, consistent with our cryo-em+MD approach.

To validate the results from both approaches and to determine the effects of each residue on OAS2 activation by RNA, we tested the effects of mutations at the predicted RNA binding residues on OAS2 activation in cells. Nearly all tested mutations evidenced similar expression levels, except for R288A which showed significantly less expression, suggesting that this mutation affects protein stability (SI Fig. 5H, I). Additionally, nearly all mutated residues decreased OAS2 activation in cells confirming that the predicted RNA binding sites are necessary for OAS2 function (Fig. 3H, I). Mutations of residues on DII (R529E R533E and K538E R544E) were chosen based on prior OAS–RNA structures (SI Fig. 5F) ^26–28,33^. As expected, mutation of these residues at the DII RNA binding site decreased OAS2 activity in cells. Notably, mutations on DI residues of interface B on the N-terminal lobe located more distantly from DII (R92, R165, D168, K169), demonstrated a significant effect on OAS2 activation (Fig. 3H, I). Their location at the opposite end of the RNA relative to DII suggests that these residues are responsible for distinguishing between short and longer RNA for activation (Fig. 3F, J). Based on the integration of cryo-em, MD simulations, and AF3 predictions, we propose that the cryo-em+MD simulations approach illustrates the RNA-induced transition from the inactive cryo-em dimer structure to the active monomeric state, while the AF3+MD predictions are most representative of the monomeric active conformation (Fig. 3J). Our results define a mechanism of OAS2 activation in which RNA must bind to both domains DII and DI to induce the conformational changes necessary for activation. The catalytically inactive DI domain functions as a molecular ruler that measures and determines the minimum RNA length required for OAS2 activation.

### OAS2 dimerization and localization to the Golgi membrane via myristoylation is required for activation

Blue-native PAGE analysis showed that the wild-type OAS2 also formed a dimer in cells as shown previously, while mutations in the zinc coordination site (C652S/H, H654S) resulted in monomeric OAS2 validating our cryo-em structure (Fig. 4A) ^34^. However, contrary to our biochemical *in vitro* findings where OAS2 C652S was hyperactive in the presence of RNA, the monomeric OAS2 mutants displayed reduced activity compared to wild-type OAS2 in HEK293T cells after poly I:C stimulation (Fig. 4B, SI Fig. 6A, B). These differences could not be explained by variations in expression levels or protein stability between OAS2 wild-type and mutant proteins (SI Fig. 6C). We hypothesized that the observed differences in activity might be explained by the mislocalization of the monomeric OAS2 mutants within cells. In BJ primary human foreskin fibroblasts (HFFs) and human lung adenocarcinoma A549 cells stimulated with IFN-α, endogenous OAS2 localized to structures marked by the Golgi marker GM130 (Fig. 4C, SI Fig. 6D, E)^57^. OAS2 was absent in cells without IFN-α stimulation or in CRISPR/Cas9-mediated OAS2 knockout cells (SI Fig. 6F, G). In A549 cells endogenous OAS2 localized to cis-Golgi, medial-Golgi, and trans-Golgi network labeled by GM130, giantin, and TGN46, respectively ^58^, but did not colocalize with the ER marker calnexin (SI Fig. 6H, I). Next, we generated A549 OAS2-KO cells reconstituted with doxycycline inducible OAS2 wild-type, OAS2 C652S and we also introduced the OAS2 G2A mutation, which disrupts a previously identified myristoylation site (SI Fig 6F) ^42^. We also confirmed that OAS2 is myristoylated in our experiments by mass spectrometry analysis (SI Fig. 6J). The OAS2 G2A myristoylation mutation disrupted OAS2 targeting to Golgi membranes rendering a diffuse cytosolic location (Fig. 4D, E). Interestingly, OAS2 C652S as well displayed a more diffuse cytosolic localization, with partial targeting to the Golgi, indicating an altered distribution compared to wild-type OAS2. These results reveal that OAS2 is a dimer localized to the Golgi(-derived) membrane and that dimerization is required for the proper targeting of OAS2 to the Golgi.

**Figure 6.**
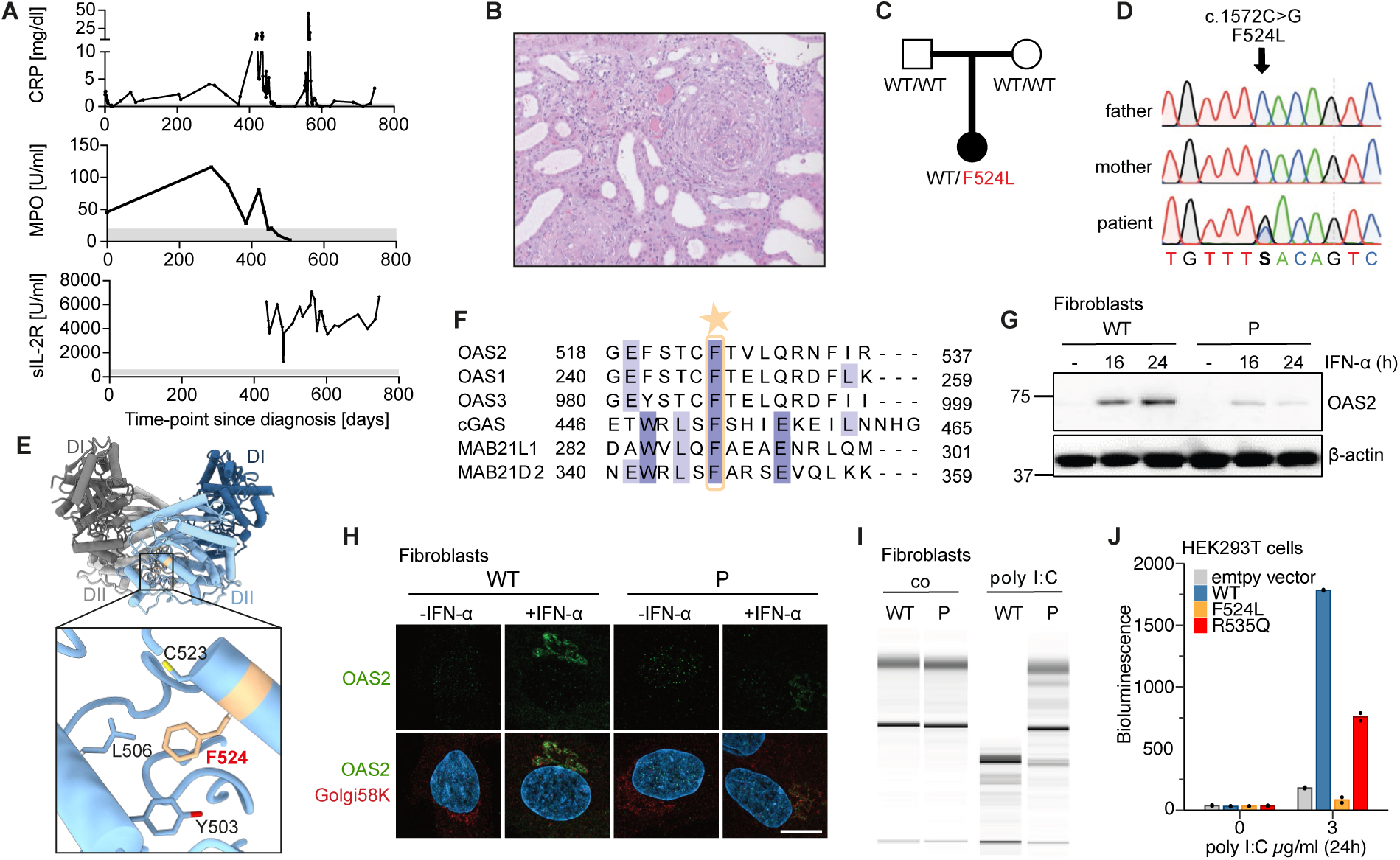
OAS2 loss of function causes immune dysregulation with ANCA vasculitis. (A) Patient’s serum levels of inflammatory markers C-reactive protein (CRP; [<0.5 mg/dl] and soluble interleukin-2 receptor (sIL-2R; [158-613 U/ml]), and antibodies against myeloperoxidase (MPO; [<20 U/ml]). Grey bars indicate normal ranges. (B) Histopathological findings of the renal biopsy showing areas of a proliferative extracapillary glomerulonephritis with cellular crescents, necrotizing vasculitis of an interlobular artery, along with tubular epithelial damage with tubular atrophy and interstitial fibrosis, as well as interstitial inflammation, consistent with ANCA-associated vasculitis. Haematoxylin and eosin stain, 20x magnification. (C) Pedigree of family with OAS2 F524L mutation. (D) Electropherograms showing *de novo* heterozygous *OAS2* variant (c.1572C>G; p.Phe524Leu) by Sanger sequencing. (E) Structure of the OAS2 dimer, with each monomer shown in grey and blue. A close-up view highlights the amino acid F524, colored in red (F) Multiple sequence alignment of human OAS proteins and selected human NTases showing high degree of conservation for F524. F524 is marked with a yellow star. (G) Expression levels of OAS2 in wild-type (WT) and patient fibroblasts either treated with IFN-α for 16 h or 24 h or left untreated. β-Actin was probed as loading control. (n=2). (H) Immunofluorescence staining for OAS2 and Golgi 58K in patient and wild-type fibroblasts. Scale bar = 10µm. (I) RNA pico-chip analysis of total RNA isolated from wild-type (WT) and patient-derived fibroblasts cell lines untreated (co) and transiently transfected with poly I:C for 24 h. (J) Analysis of OAS protein activity in cells with 2′-5′OA biosensor for OAS2 wild-type and patient mutations F524L and R535Q showing loss of activity for the F524L mutant. HEK293T were transiently transfected with different OAS2 constructs for 48h followed by treatment with poly I:C for 24h. Assay is representative of three independent experiments. Bars represent means of technical replicates (dots).

To further define how the cellular localization of OAS2 impacts its activity, we investigated the activity of OAS2 wild-type and the differently localizing OAS2 mutants C652S and G2A in HEK293T cells after stimulation with poly I:C. Unexpectedly, cytosolic OAS2 G2A showed no activity, indicating that Golgi localization is essential for OAS2 activation (Fig. 4F, SI Fig. 6K, L), explaining why OAS2 C652S is less active in cells. To further demonstrate that the dimer-to-monomer transition induced by dsRNA is essential for OAS2 activation, we aimed to generate an OAS2 mutant that remains a dimer but has reduced affinity between the monomers, thereby facilitating activation by RNA. We targeted residues S150A D153A at the dimer interface that we predicted to have a lesser impact on oligomerization compared to other zinc-coordinating residues when mutated (Fig.4G). As expected, OAS2 S150A D153A was dimeric in cells and showed higher activity at earlier time points at 4h and 8h after poly I:C stimulation than OAS2 wild-type (Fig.4H, I, SI Fig. 6M, N). Our results demonstrate that in addition to its RNA ligand, OAS2 requires both Golgi membrane targeting via myristoylation and RNA-dependent dimer-to-monomer transition for activation.

These findings align with the recent discovery that OAS1 isoform p46 is prenylated and targeted to membranes of the endomembrane system, further highlighting the importance of membrane localization in the function of OAS family members ^19,20,41,59^. To discern the localization of OAS2 and OAS1 isoform p46, we treated primary HFFs that express OAS2 and OAS1 p46 with IFN-α (SI Fig. 7A). OAS1 partially colocalized with OAS2 but also exhibited a more diffuse localization that may correspond to its known localization in the ER (SI Fig. S7B). To test how the precise membrane localization and orientation of the OAS domain relative to the membrane affects its activity we designed a series of chimeric proteins: OAS2 G2A myristoylation-deficient construct with the C-terminal prenylation motif found in OAS1 isoform p46, and OAS1 p46 ATIL prenylation-deficient construct that was fused with the N-terminal myristoylation motif from OAS2. Interestingly, myristoylated OAS1 p46 ATIL was targeted to the Golgi similar to the wild-type OAS1 p46 whereas prenylated OAS2 G2A formed cytosolic aggregates distinct from Golgi-localized wild-type OAS2 (SI Fig. 7C, D, E). The chimeric proteins did not produce any 2’-5’-OA, which may be due to reduced expression levels for e.g. OAS1 p46 chimera (SI Fig.7E, F) and to diffuse distribution of OAS1 p42 and OAS1 p46 ATIL as for OAS2 G2A. In addition, we fused either the myristoylation motif containing N-terminus or prenylation motif containing C-terminus to the cytosolic protein OAS3. Targeting OAS3 to membranes not only prevented its activation in the absence of poly I:C stimulation, but also reduced 2’-5’-OA production by OAS3 to levels similar to those observed for OAS1 p46 and OAS2 (SI Fig.7E, F). Together, our results define the cellular requirements for activation of OAS2: the localization and dimerization of OAS2 at the Golgi membrane are essential for its activation, and cytosolic RNA alone is insufficient to activate the protein in cells. These results challenge the previous dogma that OAS2 functions solely as a cytosolic RNA sensor.

### OAS2 restricts viruses replicating at the endomembrane system

Several viruses are restricted by members of the OAS protein family, with OAS3 previously playing a dominant role in cytosolic viral RNA restriction ^40,54–56,60–62^. To identify which viruses are sensed by OAS2 at the Golgi membrane and to further elucidate OAS2 activation in cells, we screened a panel of eleven GFP-reporter and non-reporter viruses for their replication in A549 OAS2 KO cells that were reconstituted to express doxycycline-induced OAS2 wild-type, and which were or were not treated with IFN-α. Live cell imaging was conducted to analyze virus replication through GFP signal detection and to assess cell death using propidium iodide staining (Fig. 5A, SI Fig. 8A, B). Our panel of viruses included: i) flaviviruses YFV and ZIKV, which have positive-sense RNA genomes and replicate in ER-derived membranes; ii) negative-sense RNA viruses VSV and bunyaviruses BUNV, LACV, and RVFV, which replicate in the cytosol; iii) negative-sense RNA viruses IAV and THOV from the Orthomyxoviridae family, which replicate in the nucleus; iv) the positive-sense RNA virus EMCV, which replicates in ER/Golgi-derived organelles and double-membrane vesicles; and v) dsDNA viruses HSV1 and VACV, which replicate in the nucleus and ER-enclosed sites in the cytosol, respectively. Of all the viruses tested, expression of the wild-type OAS2 specifically reduced virus-induced cell death only during EMCV infection (SI Fig. 8A, B), an effect that was not observed in cells expressing the OAS2 C652S or OAS2 G2A (Fig. 5B). Additionally, OAS2 expression decreased intracellular dsRNA levels in EMCV infected cells (Fig. 5E) in an RNase L-dependent manner (SI Fig. 8C) and OAS2 localized to viral dsRNA during viral infection (Fig. 5C, D).

EMCV is a cardiovirus that utilizes membranes from the endomembrane system to create viral replication organelles characterized by double-membrane vesicle (DMV) structures. Since EMCV was the only virus restricted by OAS2 in our screen, this raises the question of whether OAS2 may also restrict other viruses that utilize a similar replication strategy. Thus we selected enterovirus Rhinovirus HRV16 and coronaviruses SARS-CoV-2, OC43, and NL63, which use Golgi or ER membranes for replication and form DMVs ^63^. In HeLa H1 cells, HRV16 viral RNA levels were significantly reduced upon overexpression of OAS2 wild-type isoform p69 and to a lesser extend p71, as well as OAS1 isoform p46 (Fig. 5F, SI Fig. 8D). OAS1 p46 localizes to the ER and Golgi and was used as a positive control here as it has previously been shown to restrict Rhinovirus replication ^20^. Similarly, overexpression of OAS2 in HEK293T cells resulted only in a slight reduction in SARS-CoV-2 viral RNA levels, but significantly reduced viral RNA levels for the common cold coronaviruses NL63 and OC43 (Fig. 5G). OC43 RNA levels were also reduced upon doxycycline-inducible expression of OAS2 wild-type but not OAS2 G2A and OAS2 C652S mutants in A549 OAS2 KO cells in the absence but not presence of IFN-α pre-stimulation (SI Fig. 8E). Taken together, our results show that OAS2 has evolved to restrict viruses that exploit the endomembrane system for replication to form viral replication organelles and DMVs.

### OAS2 loss of function causes immune dysregulation with ANCA vasculitis

To explore the relevance of our findings to human physiology, we analyzed patients with early onset immune dysregulation and identified a heterozygous OAS2 variant in a child with autoinflammation and autoimmunity of unknown cause (Supplementary Data, extended case report). The girl presented with rapidly progressive glomerulonephritis at the age of 4 years. She was tested positive for perinuclear anti-neutrophil cytoplasmic antibodies (p-ANCA), antibodies to myeloperoxidase (MPO), and antinuclear antibodies (ANA) (Fig. 6A, SI Fig. 9A). A kidney biopsy showed necrotizing and crescentic glomerulonephritis, consistent with MPO-ANCA-associated vasculitis (Fig. 6B, SI Fig. 9A). Throughout the course of her illness, the patient was found to have elevated inflammatory markers (Fig. 6A, Sl Fig. 9A). Despite repeated negative tests for infections, she was treated with antibiotic therapy. She also received immunomodulatory therapy with steroid pulses, anakinra, and infliximab. However, hyperinflammation could not be controlled and the patient developed acute coronary syndrome with myocardial infarction with severe hemorrhagic complications. At the age of 6 years, the child suffered another myocardial infarction with a massive increase in inflammatory markers. A cytokine storm due to bacterial sepsis was suspected as the cause of the circulatory failure. Despite multimodal intensive care support, the child succumbed to cardiocirculatory failure.

Whole exome sequencing revealed a heterozygous variant in OAS2 (c.1572C>G, p.(Phe524Leu), NM_002535.3) in the patient, affecting a highly conserved amino acid residue of the OAS2 protein (Fig. 6C, D, SI Fig. 9B). Neither of the healthy parents carried this variant, indicating that it had arisen de novo (Fig. 6C, D). The F524L variant was not reported in the gnomAD database v4.1.0 and was predicted to be possibly damaging with a PolyPhe-2 score of 0.953. Conservation of F524 extends beyond OAS2, as this residue is highly conserved in other proteins harbouring the NTase fold (Fig. 6F) ^6^. We mapped F524 to the catalytically active domain DII of OAS2 in our cryo-em structure, where it is inserted into a hydrophobic pocket with multiple contacts to other hydrophobic residues (Fig. 6E). Other patient mutations R535Q, Q258L and V290I on OAS2, previously identified in children multisystem inflammatory syndrome (MIS-C), mapped to different sites of our cryo-em structure: i) R535 is located on a loop in DII close to the RNA binding interface, while Q258L and V290I are in close proximity located on DI (SI Fig. 9C) ^24^. These mutations were described to have similar expression as wild-type OAS2 but with altered activity. Based on our structure and the effects of previous patient mutations, we predicted that mutation of F524 to leucine would drastically disrupt protein structure as well as stability and thus cause a loss of function. Consistent with our predictions, patient fibroblasts showed significantly lower OAS2 protein expression compared to wild-type fibroblasts (Fig. 6G). After stimulation with IFN-□, OAS2 still localized to the Golgi in patient fibroblasts, although at lower levels compared to wild-type cells, reflecting heterozygosity for F524L (Fig. 6H). In agreement, activation of RNase L was significantly reduced in patient fibroblasts after treatment with poly I:C as well as in A549 OAS2 KO cells reconstituted with OAS2 F524L (Fig. 6I, SI Fig. 9D, E). Overexpression of OAS2 F524L in HEK293T showed no activity when using the biosensor for 2’-5’OA even 24 hours after stimulation with poly I:C. In contrast, the previously reported patient mutation R535Q showed only reduced activity, indicating that the F524L has a significantly stronger effect on OAS2 function. (Fig. 6J, SI Fig. 9F, G). In contrast to healthy control cells, patient fibroblasts proliferated slower and showed significantly reduced translational activity, as shown by metabolic labeling of protein synthesis (SI Fig. 9H, I, J). While treatment with IFN-□ resulted in translational shutdown in wild-type cells, it had no further effect on translation inhibition in patient cells (SI Fig. 9I, J), suggesting an integrated stress response in patient cells, possibly due to the presence of misfolded mutant OAS2 protein. Taken together, we identified a heterozygous de novo OAS2 mutation associated with a complete loss of function in a child with a severe autoinflammatory and autoimmune phenotype.

## Discussion

Our results define an activation mechanism for a multi-domain OAS protein in which OAS2 exists in an autoinhibited state as a zinc-mediated dimer, allowing it to be activated by dsRNA in a length-dependent manner. While the catalytically active domain DII of OAS2 is trapped within the dimer in an auto-repressed conformation, domain DI acts like a molecular ruler that measures the length of the RNA required for activation. We demonstrate that dimerization is important to localize OAS2 to Golgi membranes via myristoylation and that this Golgi membrane-association is required for OAS2 activation. This mechanism allows OAS2 to specifically restrict several viruses that exploit the endomembrane system for replication, such as picornaviruses, coronaviruses and rhinoviruses. Our results underscore the non-redundant role of OAS proteins in viral restriction and RNase L activation, and we further emphasise the clinical relevance of OAS2 by identifying a patient with immune dysregulation carrying a heterozygous loss-of-function mutation in OAS2.

Historically, OAS protein activation has been primarily characterized by its convergence on RNase L activation, with little understanding into the distinct antiviral roles of various OAS genes and even less clarity regarding the different isoforms of OAS genes. In this context, it has been widely described that OAS proteins have evolved to discriminate between RNA lengths, broadening the spectrum of viral RNAs to be detected during a viral infection ^47,64–68^. However, the mechanism for length discrimination by members of the OAS protein family with multiple OAS domains resulting from gene duplication and domain coupling remained unclear. The fact that certain OAS domains have lost their catalytic activity has further obscured our understanding of function of the OAS protein family. A current model is that both domains of OAS2 are required for activation, as the isolated catalytically active domain DII of OAS2 was unexpectedly found to be inactive in the presence of RNA ^32,34–36^. Our data on the dimeric structure of OAS2 explain the underlying mechanism: domain DII is held in an inactive conformation by a zinc coordination site, while DI provides the necessary trigger to release DII from the inactive conformation after RNA binding. Auto-inhibition of OAS2 by dimerization represents the first identified mechanism for RNA length discrimination in the OAS protein family and shows how OAS2 is prevented from being activated by shorter RNAs, thereby avoiding autoreactivity. This regulatory mechanism, unique to OAS2, also explains its different ligand requirements compared to OAS1 and OAS3.

The recent discovery that C-terminal prenylation of the OAS1 isoform p46 targets OAS1 to the membranes of the endomembrane system for virus detection has sparked a new perspective on investigating the non-redundant roles of OAS proteins ^19,20^. Our results showing that proper targeting of OAS2 to the Golgi by N-terminal myristoylation is required for OAS2 activation further support the notion that members of the OAS protein family have evolved not only to recognize different RNA ligands, but more importantly to restrict viral replication at different subcellular localizations. Its localization at the Golgi membrane allows OAS2 to specifically detect viruses that utilize membranes to form replication organelles with double-membrane vesicle structures and that would otherwise effectively protect their RNA from being recognized by the immune system. This feature distinguishes the antiviral repertoire of OAS2 from that of other RNA sensors, such as OAS3 which senses viruses that replicate in the cytosol ^40^. Positioning of OAS proteins at viral replication organelles, i.e. sites that are rich in viral RNA and increasing the local OAS2 concentrations, would also explain why OAS proteins can sense slight changes in dsRNA in cells despite the previously observed relatively low affinities of OAS1 and OAS2 for RNA ^35,51,66,69^. In addition, differences in virus restriction among members of the OAS family may also be attributed to evasion mechanisms that viruses have evolved^70^.

In line with recent findings that OAS1 and OAS2 restrict SARS-CoV-2, autosomal recessive deficiencies in OAS1, OAS2, or RNase L have been shown to result in multisystem inflammatory syndrome (MIS-C), a hyperinflammatory condition following SARS-CoV-2 infection ^24^. We identified a child with a de novo mutation in OAS2, F524L, who developed severe autoinflammation and autoimmunity in the context of ANCA-vasculitis. Unlike the hypomorphic OAS2 mutations associated with MIS-C, F524L causes a complete loss of function of OAS2 by destabilizing its structure. The associated phenotype of autoinflammation and autoimmunity supports the idea that OAS proteins have evolved to restrict different viruses and that these have non-redundant roles in antiviral immunity. It is tempting to speculate that the patient was infected with a common cold virus, such as coronaviruses or rhinoviruses, which are restricted by OAS2 and not usually tested for during hospitalisation. Persistent infection due to defective restriction of the virus may have been the trigger for the autoinflammatory and autoimmune phenotype. Our results reveal the first mechanisms of auto-inhibition and activation of a multi-domain OAS protein, illustrate how subcellular localization in cells is required for OAS2 to exert its antiviral function, and highlight the clinical relevance of OAS2.

## Acknowledgements

We thank all the members of the NTase laboratory for helpful comments and discussion. V6 and V6-Y312A in pCDNA4/ TO 2’-5’-oligoadenylate biosensors were kindly provided by A. Korennykh, Princeton University. We thank K. Krey for his guidance on virus infections and IncuCyte live cell imaging, as well as K. Austen for her assistance with Airyscan microscopy. We thank J.-R. Fischer and B. Ott for technical assistance, L. Kuhn for cloning and providing the OAS protein plasmids for Rhinovirus infections and G. Witte for advice on mass photometry experiments. We thank P. Ascanio and S. D′Souza for help with cloning OAS2 constructs. We are thankful to K. Hackmann, Institute for Clinical Genetics, TU Dresden, for NGS data processing, and K. Amann, Institute of Pathology, FAU Erlangen-Nürnberg, Erlangen, for renal histology. The authors gratefully acknowledge the scientific support and HPC resources provided by the Erlangen National High Performance Computing Center (NHR@FAU) of the Friedrich-Alexander-Universität Erlangen-Nürnberg (FAU) under the NHR project b119ee.

The work was funded by the German Research Foundation Emmy Noether Programm 458004906 (C.C.O.M.), CRC237 369799452/A25/A11,B11/A07/A28 (C.C.O.M., M.L.-K., A.P., L.K.), CRC369 501752319/C06 (M.L.-K.), CRC1279/A08, SP 1600/7-1, SP 1600/9-1 (K.M.J.S.), Sachbeihilfe grant KM 5/3-1 (D.K.), 52953824 (N.S., A.S.), European Research Council ERC-StG-2023 101117085 (C.C.O.M.), the German Federal Ministry of Education and Research BMBF IMMUNOMOD-01KI2014 (K.M.J.S.) and 01GM2206C (GAIN, M.L.-K.), 01GL2405H (DZKJ, M.L.-K.) I.B. is supported by a DFG Walter Benjamin Fellowship. R.S. and D.K. were supported by a Baustein Grant of the Medical Faculty, Ulm University. The Orbitrap Eclipse mass spectrometer was funded in part by the German Research Foundation (INST 95/1650-1 FUGG).

## Author Contributions

The project was conceived, and experiments designed by V.M., I.B. and C.C.O.M. V.M. prepared cryo-em samples and performed biochemical analysis with assistance from S.H. and C.C.O.M. V.M. and C.C.O.M. optimized and screened cryo-em grids. K.L. collected cryo-em data. V.M. performed structure determination and model building with assistance from K.L. and C.C.O.M. MD simulations were performed by A.S. and N.S. I.B. performed cell-based assays and fluorescence microscopy. M.H. quantified fluorescence microscopy images. I.B., D.K. and R.S. performed virus infections under supervision of C.C.O.M., A.P. and K.M.J.S. I.B. prepared samples for mass spectrometry and J.M. collected and analysed data. S.K., R.L. and I.B. performed patient-related experiments under supervision of M.L.-K. and C.C.O.M. M.L.-K. analysed genomic patient data and supervised patient-related experiments and interpreted data. K.B. and M.W. contributed patient data. J.T. and L.M.K. provided IVT RNA. The manuscript was written by V.M., I.B. and C.C.O.M. and all authors contributed to editing the manuscript and support the conclusions. The authors gratefully acknowledge the scientific support and HPC resources provided by the Erlangen National High Performance Computing Center (NHR@FAU) of the Friedrich-Alexander-Universität Erlangen-Nürnberg (FAU) under the NHR project b119ee.

## Experimental Procedures

### Protein expression and purification

Human OAS2 wild-type and OAS2 constructs were cloned into a modified pACBac1 vector with N-terminal His-tag. The plasmid was transformed into *E. coli* DH10alphaMultiBac and plated on LB-agar plates containing 7 µg/ml gentamicin, 50 µg/ml kanamycin, 12.5 µg/ml tetracyline, 0.05 mM isopropyl thiogalactoside (IPTG), and 0.1 mg / ml X-Gal. Baculovirus DNA was extracted using NucleoBond Xtra Midi Kit (Macherey-Nagel) and transfected in *Spodoptera frugiperda* Sf9 insect cells. The Baculovirus was propagated twice in Sf9 cells and used for 1:100 infection of 2 L *Trichoplusia ni* High Five cells at a concentration of 1.0 x 10^6^ cells/mL. Cells were cultured for 72 h at 27°C and harvested by centrifugation.

Cells were lysed by sonification in lysis buffer (20 mM HEPES pH 7.5, 400 mM NaCl, 10% glycerol, 30 mM imidazole, 1 mM β-mercaptoethanol, protease inhibitor (100 mM PMSF, 200 mM benzamidine, 200 µM pepstatin A, 60 µM leupeptin). To clear the lysate, sample was centrifuged at 20.000 rpm for 1h at 4°C. Cleared lysate was added to equilibrated Ni-NTA resin (Macherey-Nagel) and incubated for 2h at 4°C. The beads were washed with 1 CV with lysis buffer, 3 CV with wash buffer (20 mM HEPES pH 7.5, 1 M NaCl, 10% glycerol, 30 mM Imidazole, 1 mM β-mercaptoethanol, protease inhibitor) in batch. The last washing step of 1 CV lysis buffer was performed on gravity flow column. The protein was eluted in 10 mL fractions with elution buffer (20 mM HEPES pH 7.0, 200 mM NaCl, 10% glycerol, 300 mM Imidazole,1 mM β-mercaptoethanol, protease inhibitor). The elution was diluted 1:3 in buffer A (25 mM HEPES pH 7.0, 100 mM NaCl, 2 mM DTT) and applied to a 5 mL HiTrap Heparin HP column (Cytiva). The bound protein was eluted with 5 CV buffer B (25 mM HEPES pH 7.0, 1 M NaCl, 2 mM DTT). Fractions were pooled and loaded on a S200 Superdex 10/300 GL (Cytiva) in SEC buffer (20 mM HEPES, pH 7.0, 250 mL NaCl, 1 mM TCEP). Purified proteins were concentrated to 2 mg/mL, flash frozen in liquid nitrogen and stored at -80°C. N-terminally His-tagged human OAS3 was cloned in pACEBac1. Purification was performed as described above, but frozen in final concentration of 3 mg/mL. Human OAS1 was purified as described before ^71^. Briefly, His-OAS1 in pET-SUMO was expressed in *E. coli* Rosetta cells, induced with 0.4 mM IPTG and grown overnight at 18°C. Cells were lysed by sonification in lysis buffer (20 mM HEPES pH 7.5, 400 mM NaCl, 10% glycerol, 30 mM imidazole, 1 mM β-mercaptoethanol, protease inhibitor (100 mM PMSF, 200 mM benzamidine, 200 µM pepstatin A, 60 µM leupeptin). Cleared lysate was loaded on Ni-NTA resin (Macherey-Nagel) in gravity flow column and washed 1 CV with lysis buffer, 3 CV with wash buffer (20 mM HEPES pH 7.5, 1 M NaCl, 10% glycerol, 30 mM imidazole, 1 mM β-mercaptoethanol, protease inhibitor) and 1 CV lysis buffer. The protein was eluted in 10 mL fractions with elution buffer (20 mM HEPES pH 7.5, 200 mM NaCl, 10% Glycerol, 300 mM Imidazole,1 mM β-mercaptoethanol, protease inhibitor) and dialyzed overnight against dialysis buffer (25 mM HEPES pH 7.5, 100 mM NaCl, 2 mM β-mercaptoethanol). During dialysis, His-OAS1 was cleaved with SUMO protease 1 (in-house production). The sample was loaded on a 5 mL HiTrap Heparin HP column (Cytiva) with buffer A (25 mM HEPES pH 7.5, 100 mM NaCl, 2 mM DTT) and eluted with 5 CV buffer B (25 mM HEPES pH 7.5, 1 M NaCl, 2 mM DTT). Fractions were pooled and loaded on a HiLoad 16/600 Superdex 75 size exclusion chromatography column (Cytiva) in 20 mM HEPES pH 7.5, 250 mM NaCl, 1 mM TCEP. Purified OAS1 was pooled and stored at -80°C.

### Cryo-electron microscopy grid preparation

For cryo-em experiments, OAS2 wild type protein was diluted to 5 µM in 25 mM HEPES pH 7.5, 85 mM NaCl and 1 mM TCEP and octyl-β-glucoside was added to final concentration of 0.05%. Quantifoil Cu 200 R2/1 grids were plasma glow discharged for 7s at 20 mA (GloQube, Quorum). Grids were prepared with a Leica EM GP plunge freezer (Leica), at 10 °C and 90% humidity. 4.5 µL protein sample was applied to grids and blotted 2.0 s before vitrification in liquid ethane.

### Cryo-electron microscopy data acquisition

Data were collected on a Titan Krios G3 transmission electron microscope (Thermo Fisher Scientific) used at 300 kV and a Gatan K2 Summit direct electron detector operated in counting mode and an energy filter. Data acquisition was automated with EPU software package (Thermo Fisher Scientific). 5708 movies were collected with defocus values ranging from -1.4 and -2.9 µM, pixel size of 1.059 Å and total electron dose of 50 e^-^Å^-2^. A representative micrograph is shown in SI. Fig. 1B.

### Cryo-electron microscopy data processing

Micrograph movies were motion corrected and dose-weighted using MotionCor2 ^72^. The CTF parameters of the datasets were estimated with CTFFIND4 ^73^. All cryo-em data processing steps were performed using cryoSPARC 4.5.3. The exact processing scheme, data collection and refinement statistics are depicted in Supplementary Fig. 1C and Table S1. 500 particles were manually picked on 5708 micrographs, extracted using 180 px box size and sorted by 2D classes. Topaz train was used for further, refined particle picking. The resulting 1729799 particles were used to generate an 3D ab-initio reconstruction and subsequently subjected to heterogeneous refinement. The best classes were selected for local CTF refinement and homogeneous refinement. The selected particles were then 3D classified. The best 3D classes were used for final homogeneous, non-uniform refinement and local refinement to obtain a map with a final resolution of 3.30 Å, calculated with the gold-standard Fourier shell correlation criterion (FSC = 0.143).

### Model building and Figure preparation

Predicted structure of OAS2 from Alphafold1 (AF-P29728-F1) was rigid-body fitted to the electron density in UCSF ChimeraX ^74^. Model building and iterative refinement were performed in Coot 0.9.8.6 with secondary structure restrains and Phenix 1.20. Figures were prepared using UCSF ChimeraX ^75^.

### In-vitro OAS activity assay

The activity of OAS was monitored by indirect colorimetric assay, measuring the byproduct pyrophosphate of the oligoadenylate synthesis ^71^. OAS protein was incubated with dsRNA (concentrations indicated for each experiment) in reaction buffer (25 mM HEPES pH 7.0, 7 mM MgCl_2_, 10 mM NaCl, 1 mM TCEP, 2 mM ATP) in a total reaction volume of 150 µL at 37°C for 120 min. To quench the reaction, 10 µL aliquots were added to 2.5 µL 0.25 M EDTA, pH 8.0 in 96 well plate at 0, 5, 10, 20, 40, 60, 80, 100 and 120 min. 10 µL 2.5 % (w/v) ammonium molybdate in 2.5 M H_2_SO_4_ and 10 µL 0.5 M β-mercaptoethanol was added to each well and the final volume were adjusted to 100 µL. The produced molybdophosphoric acid was reduced to molybdenum blue complex that was measured at absorbance at 580 nm. Data were compared with pyrophosphate standard and analyzed using non-linear regression in GraphPad Prism 8.

### EMSA (electromobility shift assay)

The affinity of OAS proteins to dsRNA was determined by electromobility shift assays (EMSA). RNA (200 nM) was titrated to increasing concentrations of OAS protein (0-14 µM) in binding buffer (5 mM MgCl_2_, 25 mM HEPES pH 7.0, 10 mM NaCl, 1 mM TCEP). The reaction was incubated for 10 min at RT and mixed with glycerol to an end concentration of 10%. Native TB gels (8%) were pre-run for 30 min. Samples were loaded and electrophoresis was performed at 4°C for 50 min at 200 V in 0.5x TB running buffer. Gels were stained in GelRed (Biotium) for 30 min at RT.

### In-vitro Transcription

3p-dsRNA-hp was synthesized by *in vitro* transcription (IVT) from a dsDNA template (Metabion, Munich) using the HiScribe^TM^ T7 Quick HighYield RNA Synthesis Kit (New England Biolabs GmbH, Frankfurt, Germany) according to the manufacturer’s protocol for small RNAs. DNase I (New England Biolabs GmbH, Frankfurt, Germany) was added to the IVT reaction for removal of the template DNA for 30 min at 37°C. A two-step purification protocol was used to achieve high purity RNA transcripts. First, RNA transcripts were separated from residual enzymes by phenol-chloroform extraction followed by ethanol precipitation. Second, the product was further separated from IVT by-products using a 12% TBE-urea polyacrylamide gel electrophoresis (PAGE). The RNA was visualized via UV shadowing on a silica-coated TLC plate, excised and crushed into small gel pieces by centrifuging it at maximum speed through a perforated 0.5 ml tube into a 1.5 ml collection tube. 3p-RNA-hp was eluted from the gel with 0.5 M ammonium acetate by shaking overnight at 16°C, precipitated with ethanol (3 volumes) and dissolved in nuclease-free water. Products were quality controlled by mass spectrometry (ESI-TOF) at Metabion, Munich.

### Mass photometry

Mass photometry measurements of OAS proteins were performed with Two MP mass photometer (Refeyn). Samples were diluted to 50 nM in filtered buffer (25 mM HEPES pH 7.0, 50 mM NaCl, 1 mM TCEP) and movies were recorded for 60s. Data were analyzed using AquireMP software (Refeyn).

### MD simulations

We used all-atom MD simulations to characterize the conformational dynamics and to predict the RNA-binding site on OAS2. OAS2 was simulated as monomer and dimer. Both systems were simulated with and without dsRNA with 44 bp. All systems were prepared with a bulk salt concentration of 100 mM NaCl. The force fields parameters were taken from the AMBER ff19SB force field ^76^, RNA force-field parameters from the AMBER99sb*ILDN-parmbsc0-χ-OL3 force field, OPC water ^77^ and ion force fields from ^78,79^. After equilibration, the production runs were done in the NPT ensemble with velocity-rescaling thermostat ^80^ with a time constant of 1 ps^-1^ and Parinello-Rahman barostat ^81^ with a coupling constant of 5.0 ps were used. The OAS2 monomer and dimer were simulated for 100 ns each at physiological conditions. Configuration sampling simulations of the dsRNA-OAS2 complexes at 400 K were followed by contact analysis and consecutive simulations at physiological temperature and pressure. More than 2.8 μs were used to identify key residues involved in RNA binding.

In addition, the protonation state of the zinc binding site was carefully adjusted to correctly account for the zinc binding which is known to play a crucial role for the stability of the OAS2-Zn^2+^ complex. Further information on the MD simulations is provided in the supporting information.

### Cell culture

HEK293T, A549, Hela H1 and BJ hTERT^+^ HFF cell lines and BJ primary HFF cells were cultured in DMEM supplemented with 10% FBS at 37°C in 5% CO_2_. HEK293T, A549 and Hela H1 cells were passaged at a dilution 1:10 by washing with PBS and detached with 0.25% trypsin. BJ fibroblasts were passaged at a dilution 1:5 by washing with PBS and detached with 0.25% trypsin. Fibroblasts from OAS2 patient and healthy controls were derived from skin biopsies. Passage-matched primary human fibroblasts (passages 4 to 20) were cultured in Dulbecco’s modified Eagle’s medium (DMEM, Sigma, D5671) supplemented with 2 mM L-glutamine, 1x antibiotics/antimycotics, 1x non-essential amino acids (NEAA), and 10% fetal bovine serum (FBS) at 37 °C and 5% CO_2_. All cell lines were regularly tested negative for mycoplasma contamination using TaKaRA PCR Mycoplasma Detection Set (TaKaRa).

### Generation of cell lines

Two different gRNA sequences for OAS2 and non-targeting control (NTC) knockout were cloned in pLentiCRISPRv2 vector (Addgene) with hygromycin resistance. OAS2 guide RNA (gRNA) sequences were designed using the Synthego (https://design.synthego.com) design tool. Non-targeting control sequences were obtained from the GeCKO v2.0 library ^82^. Primers encoding target guide RNA sequences were annealed and used for ligation with linearized vector from restriction digest with BsmbI enzyme. Primer sequences containing overhangs for ligation are listed in the Table 1. Successful cloning was verified by sequencing and target vectors were further used for lentivirus generation.

**Table 1:**
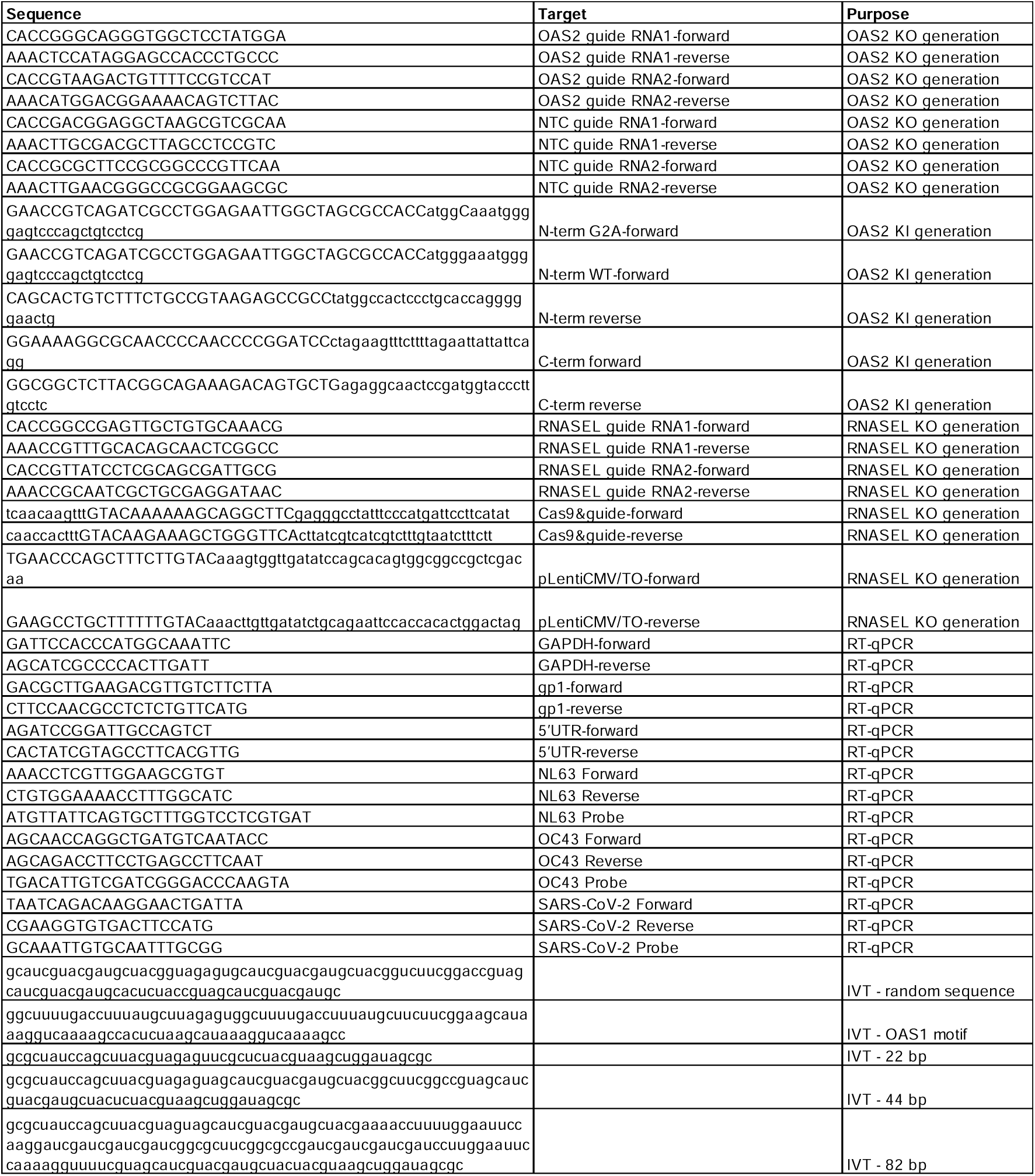
Oligonucleotides used in the study for cell line generation, RT-qPCR and IVT.

For generation of doxycycline-inducible OAS2 KI cell lines, OAS2 p71 N- and C-terminal fragments were amplified from respective OAS2 WT, OAS2 G2A, OAS2 C652S or OAS2 F524L constructs in pCDNA4 vector using primers listed in the Table 1. Two fragments from N-terminus and C-terminus were amplified from OAS2 gene to introduce silent mutations and mismatches in gRNA binding site and introduce overlaps with target vector pLIX-403 for Gibson assembly ligation. Target vector pLIX-403 was linearized using NheI and BamHI restriction enzymes. Inserts from OAS2 N- and C-termini to generate respective WT, G2A, C652S, G2A C652S or F524L constructs were used for Gibson assembly with linearized pLIX-403 vector and transformed into STBL3 *E.coli* strain.

For generation of RNASEL KO monoclonal HEK293T cell lines by transient transfection, two guide RNAs for RNASEL and NTC KO were first cloned into pLentiCRISPRv2 vector with hygromycin resistance as described above for OAS2 gRNA cloning. RNASEL gRNA sequences were designed using CHOPCHOP web tool and are listed in the Table 1. Afterwards fragment encoding gRNA and Cas9 in pLentiCRISPRv2 vector and pLentiCMV/TO (zeocin resistance) vector were amplified by PCR to generate overlaps for Gibson assembly primers listed in the Table 1. Fragments were ligated using Gibson assembly and the resulting vector was used for cell line generation.

Lentiviral transductions were used to generate A549 OAS2 KO cell lines reconstituted with doxycycline-inducible OAS2 constructs. For knockout generation lentiviral particles were generated by transfecting HEK293T cells with psPAX2 and pMD2.G-VSV-G packaging plasmids together with pLentiCRISPRv2 vector with hygromycin resistance encoding *S. pyogenes* CRISPR-Cas9 and two different gRNA sequences. Lentiviral particles were harvested 48h post transfection and used to infect A549 cells followed by selection with 500 µg/ml hygromycin one day post infection. Selection was terminated once there were no more viable control untransduced cells with hygromycin selection.

For reconstitution of OAS2 lentiviral particles were generated by transfecting HEK293T cells with psPAX2 and pMD2.G-VSV-G packaging plasmids together with transgene in pLIX_403 vector with puromycin resistance. Lentiviral particles were harvested 48h post transfection and used to infect A549 OAS2 KO cells followed by selection with 1.5 µg/ml puromycin one day post infection. Selection was terminated once there were no more viable control untransduced cells with selection. OAS2 KO and reconstitution was validated by WB analysis.

For monoclonal RNASEL KO cell line generation in HEK293T cells, cells were transfected with two different guides for RNASEL or NTC and *S. pyogenes* CRISPR-Cas9 in pLenti CMV/TO vector with zeocin resistance. 48h after transfection cells were selected with 400 µg/ml zeocin for 5-7 days until control transfected cells were not viable. Afterwards single cells were seeded into 96 well plate and grown for 7-10 days until formation of single colonies visible by bright-field microscopy. Single colonies were selected and expanded and KO was validated by WB analysis.

### Sanger sequencing

Genomic DNA was extracted from blood using the QIAamp DNA Blood Mini Kit (QIAGEN). Genomic DNA flanking the *OAS2* (ENST00000392583_7) mutation was amplified by polymerase chain reaction (PCR) using gene-specific primers [Eurofins MWG Operon; OAS2-for: GAGATGCTCCCTGTGTCTTAGA and OAS2-rev: CCCCATGGTCAGAACAAAGAC] and sequenced in both directions using the service from Eurofins Genomics. Data were analyzed using the Snapgene software.

### RNase L-split luciferase assay using 2′-5′ oligoadenylate (OA) biosensor

Measurements of in vivo 2’-5’OA synthesis was performed as previously described by ^53^ using 2′-5′OA biosensor V6 (WT) and V6-Y312A variant lacking 2’-5’OA binding capacity. For the assay, 3x10^4 HEK293T cells were seeded into flat-bottom, white 96-well plates with two wells/ technical replicates per condition. The following day cells were transfected with 75ng V6 or V6-312A plasmid and 75ng OAS expression constructs using 0.4 µl Lipofectamine 2000 (Thermo Fisher Scientific). Forty-eight hours after transfection, cells were transfected with poly(rI:rC) dsRNA (1 and 3 µg/ml) using Lipofectamine 2000 and treated with 100 µM D-luciferin ethyl ester (Biomol). Luminescence was measured in 2h intervals over 8 hours and at 20-24h using TECAN plate reader. After the last measurement, cells were harvested and used for WB analysis to validate equal protein expression.

### Western Blot analysis

For monitoring induction of OAS protein expression, cells were stimulated for 24h (unless stated differently in the figure legend) with 1 µg/ml doxycycline for A549 OAS2 KO OAS2 KI cells lines or 500 or 1000U/ml IFN-α to induce endogenous OAS proteins in A549 or BJ primary and hTert^+^ HFF cell lines. After stimulation cells were washed with PBS and harvested for Western Blot analysis. Cells were lysed in NP-40 lysis buffer (50mM Tris-HCl pH 7.5, 150mM NaCl, 1% NP-40, 5mM EDTA) supplemented with 1X Complete protease inhibitor (Sigma-Aldrich) for 20-30min on ice. Soluble fraction was separated by centrifugation for 10min, 21’000xg, 4°C, mixed with 1x Laemmli sample buffer and boiled for 5-10min at 95°C. Protein was resolved by 10% SDS-PAGE and transferred to 0.45 µm PVDF membrane. Membranes were blocked in 5% non-fat dry milk PBS 0.1% Tween-20 and incubated with the following primary antibodies: anti-OAS2 (Proteintech, 19279-1-AP), anti-OAS1 (Cell Signaling, 14498S), anti-OAS3 (Proteintech, 21915-1-AP), anti-GAPDH (Santa cruz, sc-47724), anti-FLAG (Sigma Aldrich, A8592), anti-RNase L (Santa cruz, sc-74405), anti OAS1 (Proteintech, 14955-1-AP; in Hela H1 cells). Afterwards membranes were probed with species IgG-specific HRP-conjugated secondary antibodies goat anti-rabbit IgG (Dako, P0448) or horse anti-mouse IgG (Cell Signaling, 7076) or Infrared Dye coupled secondary antibodies for Hela H1 cells. Immunoblots with HRP signal were developed with the SuperSignal West Femto kit (Thermo Fisher Scientific) and imaged with the Bio - Rad ChemiDoc Imaging System. Image Studio was used for capturing images with Infrared Dye-coupled secondary antibodies.

For interferon-mediated OAS2 induction, wild-type (WT) and patient-derived (P) fibroblasts were stimulated with IFN-α (500 IU/ml) for 16 h or 24 h before cell lysis in RIPA buffer (50 mM Tris-HCl, pH 7.4, 150 mM NaCl, 1 mM EDTA, 1% Triton X-100, 1 mM Na3VO4 and 20 mM NaF) supplemented with 2 U/ml DNase I (Invitrogen), cOmplete protease inhibitor cocktail (Roche), and PhosSTOP (Roche). Hela H1 cells were lysed on ice with transmembrane lysis buffer (50 mM HEPES pH 7.4, 150 mM NaCl, 1% Triton X-100, 5 mM EDTA) supplemented with 1:500 protease inhibitor.

### BN-PAGE analysis

BN-PAGE was performed using NativePAGE Novex 4-16% Bis-Tris Gel System (Thermo Fisher Scientific) following manufacturer’s instructions. 1.5x10^5 HEK293T cells were seeded per well in 24 well plates and the following day cells were transfected with 330ng plasmid DNA using 1ug PEI transfection reagent. 24-48h post transfection cells were harvested and lysed in 0.5% digitonin, 1x BN-PAGE sample buffer (50mM BisTris, 6N HCl, 50mM NaCl, 10% w/v Glycerol, 0.001% Ponceau S, pH 7.2) supplemented with 1xComplete protease inhibitor (Sigma-Aldrich) on ice for 20-30min. Soluble fraction was separated by centrifugation for 30min, 21’000xg, 4°C. Before loading on the BN-PAGE gel lysates were supplemented with 0.125% Coomasie G-250. Purified recombinant protein for BN-PAGE was diluted in 25mM HEPES, pH 7.0, 1mM TCEP and mixed with 1x BN-PAGE sample buffer supplemented with 1xComplete protease inhibitor (Sigma-Aldrich) and mixed with 0.125% Coomasie G-250 before loading on the gel. Proteins were separated by running on ice at 150V for 60min in Anode buffer (50mM BisTris, 50mM Tricine, pH 6.8) in the outer chamber and in Dark cathode buffer (Anode buffer supplemented with 0.02% Coomasie G-250) in the inner chamber until the dye front reaches 1/3^rd^ of the gel. Afterwards the gel was run at 250V for 60min in Light cathode buffer (Anode buffer supplemented with 0.002% Coomasie G-250) in the inner chamber until dye front reached the end of the gel gel. Protein was transferred to 0.45 µm PVDF membrane in 25mM Trizma base, 150mM Glycine, pH 8.3, 20% Methanol for 1h at 100V and fixed by incubation with 8% acetic acid for 15min at RT. Blocking and antibody incubations were performed as described in the Western Blot analysis.

### Immunofluorescence staining and imaging

For imaging of A549 cells, 3x10^4 cells were seeded in 8-well x-well cell culture chambers (Sarstedt) and following day cells were stimulated with 1υg/ml doxycycline or 500U/ml IFN-α for 24h. For imaging of EMCV dsRNA and OAS2 in A549 cells, cells were stimulated with 1ug/ml doxycycline for 24h and infected with EMCV for 24h. BJ primary and hTert^+^ fibroblasts were seeded at density 8000 cells per chamber and stimulated with 1000U/ml with IFN-α for 24h. HEK293T cells were seeded at the density 6x10^4 cells per chamber and transfected with 330ng plasmid DNA using PEI transfection reagent.

Cells were washed three times with PBS and fixed with 4% paraformaldehyde in PBS for 15-20min at RT followed by three washes with PBS. Cells were permeabilized with 0.1% TRITON-X-100 in PBS for 10min at RT, washed three times with PBS and blocked in 2% BSA in PBS, 0.1% Tween-20 for 30min at RT. Immunostaining was performed using primary antibodies: anti-OAS2 (R&D Systems, MAB1925-SP, 1:100), anti-OAS2 (Proteintech, 19279-1-AP,1:250), anti-GM130 (Thermo Fisher Scientific, 703794, 1:100), anti-OAS1 (Cell Signaling, 14498S, 1:100), anti-GM130 (Proteintech, 66662-1-Ig, 1:500), anti-calnexin (R&D Systems, NB100-1965SS, 1:200), anti-giantin (R&D Systems, NBP1-91937, 1:200), anti-TGN46 (R&D Systems, NBP1-49643SS), anti-Golgi 58K (1:200) and anti-dsRNA J2 (SCICONS, 10010500). Antibody mixes were diluted in 2% BSA in PBS, 0.1% Tween-20 and incubated with cells for 1h at RT. Cells were washed three times with PBS and incubated with fluorophore-conjugated secondary antibodies: anti-rabbit AF488 (Cell Signaling, 4412S, 1:1000) and anti-mouse AF647 (Thermo Fisher Scientific, A31571, 1:1000) in 2% BSA in PBS, 0.1% Tween-20 together with DAPI (1:1000) for 1h at RT in the dark. Cells were washed three times with PBS and mounted using ProLong Gold antifade mountant (Thermo Fisher Scientific). Super-resolution Images were acquired using a Zeiss LSM 900 (Zeiss, Oberkochen, Germany) with an Airyscan 2 detector system using a Plan-APOCHROMAT 63×/1.4 Oil DIC Objective and 40×/1.2 Oil DIC Objective. Image processing and brightness adjustments were made using the ImageJ software version 1.54. Confocal images were acquired using an Olympus FV10i microscope (Olympus, Tokyo, Japan) with a 60×/1.2 water immersion objective.

Primary OAS2 F524L and control fibroblasts were seeded on round 12 mm coverslips in 24-well plates at a density of 15,000 cells per well and incubated overnight. Cells were fixed with 4% formaldehyde for 20 min and permeabilized with 0.25% Triton X-100 in PBS for 10 min at 21 °C. After blocking with 3% BSA/ 0.3% Triton X-100 in PBS for 1h at 4 °C, cells were incubated with the appropriate primary antibodies diluted in at 4 °C for 16h. Afterwards cells were incubated with corresponding Alexa Fluor-labeled secondary antibodies (Invitrogen, Alexa Fluor 488 goat anti-rabbit IgG, A11008; Alexa Fluor 594 goat anti-mouse IgG, A11032) diluted in 1% BSA/ 0.3% Triton X-100 containing PBS for 1 h in the dark at 21 °C. Cells were washed three times with PBS before mounting with VectaShield containing DAPI. Fluorescence images were captured using a LSM780 or a LSM980 confocal microscope (Zeiss).

For quantification of OAS2 colocalization with ER and golgi markers and dsRNA, colocalization analysis was conducted with the Huygens Professional 19.10 software, employing the “Huygens Colocalization Analyzer” and the Costes method to calculate Pearson correlation coefficients.

### Live cell imaging of viral replication

For virus screen stable A549 OAS2 KO OAS2 WT KI cells were seeded at density 4000 cells/ well in 96 well plates in DMEM, 10% FBS and 1:2000 propidium iodide (1mg/ml stock) with two wells/ technical replicates per condition. The following day cells were stimulated with 1µg/ml doxycycline, 10U/ml IFN-α or a combination of both or media alone for control. 24h after stimulation cells were infected with a set of 11 viruses using two different MOIs for each virus. Viruses were kindly provided by Prof. Andreas Pichlmair and included 1) GFP reporter viruses: IAV SC35M PB2 2A GFP (MOI 1, 0.1), YFV YF-17D-Venus (MOI 0.85, 0.1), RVFV-GFP (MOI 1, 0.1), VACV-V300-GFP (MOI 1, 0.1), HSV1(17+)Lox-GFP (MOI 1, 0.1) and VSV-GFP-Indiana (MOI 0.1, 0.01); 2) non-reporter viruses: LACV (MOI 1, 0.1), ZIKV (MOI 1, BUNV (MOI 1, 0.1), THOV-ML^-^ (MOI 1, 0.1) and EMCV (MOI 0.6, 0.06). Phase (cell confluence), red fluorescence intensity from propodium iodide and green fluorescence intensity for reporter viruses were measured every 3h over 72h post infection at 37°C, 5% CO2 using IncuCyte S5 fluorescence light microscopy screening platform (Sartorius). The virus replication for reporter viruses was quantified as integrated green fluorescence intensity per image normalized to cell confluence per image. Cell death was quantified as red fluorescence area per image normalized to cell confluence per image. Data were analysed using IncuCyte S5 Software (Sartorius).

For EMCV infection of stable A549 OAS2 KO cells with OAS2 WT, OAS2 G2A, OAS2 C652S and OAS2 G2A C652S KIs, cells were treated and analysed as described above.

### EMCV, Rhino- and coronavirus infections and viral RNA detection

For analysis of intracellular EMCV RNA levels in A549 OAS2 KO KI WT cells 4.5x10^4 cells/ well were seeded in 12 well plate. The following day cells were treated with 1 µg/ml doxycycline or 10U/ml IFN-α for 24h. Afterwards cells were infected with EMCV MOI 0.3 for 24h. Cells were then washed twice with PBS and cell pellets harvested for RNA isolation. For analysis of intracellular EMCV RNA levels in HEK293T cells 1.5x10^5 cells/ well were seeded in 24 well plate. The following day cells were transfected with 330ng target plasmid using 1 µl PEI (1mg/ml stock). Afterwards cells were infected with EMCV MOI 1 for 24h. Cells were then washed twice with PBS and cell pellets harvested for RNA isolation. For analysis of EMCV RNA levels total RNA was extracted using the NucleoSpin RNA Plus kit (Macherey-Nagel) according to the manufacturers’ protocol. Total RNA was used for reverse transcription with PrimeScript RT reagent Kit with gDNA Eraser (TaKaRa) according to the manufacturers’ instructions. Relative transcript quantification was obtained by qPCR with the transcript-specific primers (Dataset EV6) and PowerUp SYBR Green master mix (Thermo Fisher) on a QuantStudio3 Real-time PCR system (Applied Biosystems). The oligonucleotides used for the analysis are listed in Table 1. Ct values were obtained using QuantStudio Design and Analysis software and averaged across technical replicates. The transcript levels were normalized to the levels of a housekeeping gene GAPDH.

For HCoV-OC43 (ATCC Cat# CR-1558TM) and NL63 (kindly provided by Lia van der Hoek) propagation, Huh-7 or LLC-MK2 cells were inoculated with a MOI of 0.1 in Dulbecco modified Eagle medium supplemented with 2% FCS. Cells were incubated at 33°C. SARS-CoV-2 (strain Netherlands/01/NL/2020, European Virus Archive, Cat# 010V 03903) was propagated on Vero E6 infected at an MOI of 0.003 in serum-free medium. For infection of HEK293T cells with coronaviruses HEK293T cells were seeded at 3*10^5 cells/ml in 24-well plates and transfected with 50ng of hACE2 and 50ng of pCG plasmid expressing GFP (control) or OAS2 p69. After 24h cells were infected with 0.1 MOI of SARS-CoV-2, NL63 or OC43. NL63 and OC43 infected cells were incubated at 33°C and SARS-CoV-2 infected cells were incubated at 37°C. The following day cells were washed in PBS and fresh medium was added. Supernatants and cells were harvested 3 days post infection and centrifuged at 3000rpm for 3min to remove cell debris.

For analysis of OC43 RNA levels in supernatants, A549 cells were seeded at density 2x10^5 cells /ml into 12-well plates in DMEM, 10% FCS, 1µg/ml doxycycline in the presence or absence of 500U/ml of IFN-β. The following day cells were infected with OC43 at MOI 0.1 and incubated at 33°C. The following day cells were washed with PBS to remove input virus. Supernatants and cells were harvested 3 days post infection and used for measurement of viral copy numbers in the supernatants by RT-qPCR.

RNA from supernatant of seasonal coronaviruses-infected cells was isolated using QIAamp Viral RNA Kits (QIAGEN) according to manufacturer’s instructions. RT-qPCR was performed with TaqMan Fast Virus 1-step master mix (Thermo Fisher, catalog no. 4444436) and OneStepPlus real-time PCR system (96-well format). Custom primers and probes (5’ FAM-TAMRA 3’) were ordered from Biomers.net and are listed in the Table 1. Synthetic SARS-CoV-2 RNA (Twist Bioscience, catalog no. 102024) or linearized plasmids encoding OC43 or NL63 N gene fragments were used as a quantitative standard to obtain viral copy numbers. All reactions were run in triplicates.

Hela-H1 cells (passage 11) were transiently transfected using lipofectamine 3000 (Thermo Fischer Scientific) with plasmid containing gene of interest or empty vector according to manufacturer’s recommendations. 24 h post-transfection cells were infected with 0.1 MOI Human Rhino Virus 16 (HRV16) purchased from ATCC, and cultured at 33 °C for 5 more days until cytopathic effects were observed. The supernatant from the infected cell cultures were collected 5 days post infection and the RNA was isolated using Viral RNA mini kit (Qiagen) according to manufacturer’s instructions. RT-qPCR was performed using TaqMan Fast Virus 1-Step Master Mix and a OneStepPlus Real-Time PCR System (96-well format, fast mode). The primer-probes for microbe detection was used (Thermo Fisher Scientific).

### Protein synthesis assay

For the detection of protein synthesis using fluorescence microscopy, fibroblasts were seeded at 15,000 cells per well on coverslips in a 24-well plate and incubated overnight at 37 °C and 5% CO_2_. The next day, cells were left untreated or stimulated with IFN-□ (500 IU/ml) for 23 h. Then, the Click-iT® Plus OPP Protein Synthesis Assay was performed accordingly to the manufacturer’s protocol. Briefly, cells were incubated with 20 μM O-propargyl-puromycin (OPP) for 1 h before fixed with 4% PFA for 15 min at room temperature. After permeabilisation for 15 min with 0.5% Triton X-100 in PBS, fluorescence labeling of the incorporated OPP was carried out by the Click-iT® reaction in the dark. After staining the DNA, cells were mounted and imaged on a Zeiss LSM980.

### Proliferation assay

Growth curve analysis of fibroblasts was performed using the Incucyte Live-Cell Analysis System (Incucyte 2020B, Sartorius). Briefly, fibroblasts were seeded at 5,000 cells per 96-well and incubated for 24 h at 37 °C and 5% CO_2_. Then, NucLight Rapid Red Cell Dye (1:1000) was prepared in DMEM and 100 µl was added per well. Phase-contrast and red channel images were taken every three hours, first scanning taking place at 24 h after adding the dye. Cell number was determined afterwards using the Analyze Particle plugin in ImageJ in individual images.

### RNA degradation assay

For analysis of OAS2 activity, fibroblasts and reconstituted A549 cells were stimulated with 2 µg/ml poly(I:C) high molecular weight (HMW) (Invivogen, tlrl-pic) for 24 h. A549 OAS2 knockout cells were treated with doxycycline (1 µg/ml) for 24 h to induce OAS2 expression. Afterwards total RNA was isolated from cells using ReliaPrep™ RNA Miniprep Systems (Promega). RNA degradation was analyzed on RNA Pico chips using Agilent Bioanalyzer 2100 (Agilent).

### Immunoprecipitation and mass spectrometry

For immunoprecipitation (IP) of OAS2, 13*10^6 cells were seeded in 15cm dish per IP and transfected with 23.4 µg plasmid DNA using 70ul PEI (1mg/ml). Cells were harvested 24-48h post transfection and lysed in 50mM Tris-HCl pH 7.5, 150mM NaCl, 1% NP-40, 5mM EDTA supplemented with 1xComplete protease inhibitor for 30min on ice. After centrifugation for 30min, at 21’000 x g 4°C, cell lysates were incubated with 4ug anti-OAS2 antibody (R&D Systems, MAB1925) and 25 µl bed volume Protein G Sepharose Beads (GE Healthcare) for 2h at 4°C. Beads were centrifuged for 12’000xg, 20sec washed once with PBS and incubated either with PBS or 100mM NEM in PBS for 4h at 4°C. Beads were washed 3-5 times in PBS, 1% NP-40, 5mM EDTA followed by one wash with PBS. Sample was eluted by heating for 10min at 70°C in 1xLithium Sample Buffer (Thermo Fisher Scientific).

In - gel trypsin digestion was performed according to standard procedures (Shevchenko et al., 2006) ^83^. Briefly, samples were run on a Nu-PAGE™ 4%–12% Bis-Tris protein gel (Thermo Fisher Scientific) for about 1 cm. Subsequently, the still not size-separated single protein band per sample was cut out, reduced (50mM dithiothreitol), alkylated (55mm chloroacetamide) and digested overnight with trypsin. The peptides obtained were dried to completeness and resuspended in 0.1% formic acid prior to liquid chromatography tandem mass spectrometry (LC-MS/MS) analysis on a 42-cm column (inner diameter: 75 microns; packed with ReproSil-Pur C18-AQ 1.9-micron beads, Dr. Maisch GmbH) coupled to an Orbitrap Eclipse mass spectrometer (Thermo Fisher Scientific). Samples were measured over a 50 min linear gradient in data dependent acquisition (DDA) and positive ionization mode followed by a targeted Parallel Reaction Monitoring (PRM) measurement for selected OAS2 peptides. Raw mass spectrometry data were processed using the MaxQuant software (version 2.0.4.0) with its built-in search engine, Andromeda ^84^. Spectra were searched against the UniProtKB database (Human, UP000005640, 75,777 entries downloaded on 01.2021). Enzyme specificity was set to trypsin, allowing for 2 missed cleavages, and search performed for glycine myristoylation (+210.19836) as a variable modification. Identifications were adjusted to 1% false discovery rate (FDR) at protein and peptide levels. Skyline (v.24.1) ^85^ was used for spectral library generation and spectrum export based on MaxQuant msms.txt output files.

### Statistical analysis

Statistical significance was calculated as indicated in Figure Legends and represented as ∗ p < 0.05, ∗∗ p < 0.01, ∗∗∗ p < 0.001, and ∗∗∗∗ p < 0.0001 using GraphPad Prism 10.3.1. and RStudio.

### Accession Numbers

The electron density reconstruction and final model were deposited at the Electron Microscopy Data Bank (EMDB) with accession code EMD-51786, and the Protein Data Bank (PDB) with accession code 9H1Z.

## Supplemental Methods Simulation setup

Using AmberTools21 ^88^ the protein topology and force field files were created based on the OAS2 cryo-em structure using the latest AMBER protein force field ff19SB ^76^. The AMBER99sb*ILDN-parmbsc0-χ-OL3 force field was used for RNA, which builds on the AMBER99 force field ^89^ and includes additional corrections to the nucleic acid backbone behavior ^90^ and glycosidic torsion ^91^.

The unstructured, extended terminal regions of the protein were removed from the cryo-em, structure, to reduce the computational costs since they are unlikely to affects dsRNA binding. Initial simulations confirmed the integrity and stability of the cryo-em structure. For MD simulations of the OAS2 monomer, half of the experimental dimer structure was used as starting point. All simulations employed the OPC water model ^77^. Na^+^ and Cl^-^ parameters were originally parameterized for TIP3P ^78^ but were used in combination with OPC water after ensuring that solvation free energies and activity derivatives were in close agreement with experiments. For the centrally coordinated Zn^2+^ ion specifically for OPC optimized parameters were used ^79^.

The protonation states of the cysteine and histidine residues, which form the Zinc coordination site essential for OAS2 dimer stability, were modified to maintain stable coordination with the Zn²⁺ ion. Preliminary simulations indicated that the standard protonation states at physiological pH result in the dissociation of the Zinc cation from the coordination site. This unphysiological behavior is likely caused by changes of the protonation state in the presence of a metal ion in the binding site. Such changes are neglected by the standard Amber tools. To provide a more realistic description of the metal binding site, the sulfur atoms of the cysteine residues were deprotonated and for the histidine residues the protonation site was shifted from the ε to the δ position.

The 44 bp dsRNA (sequence: GCGCUAUCCAGCUUACGUAGAGUAGCAUCGUACGAUGCUACGGC) for the simulations of the complex was generated using the *fd_helix.c* function (David A. Case 2018, https://casegroup.rutgers.edu/fd_helix.c). Using PyMOL [The PyMOL Molecular Graphics System, Version 3.0.3 Schrödinger, LLC.], the dsRNA structure was aligned to the 18 bp piece of dsRNA bound to the protein, as resolved in thee cryo-em structure. The initial structures were solvated, neutralized and further Na^+^ and Cl^-^ ions were added to achieve a bulk concentration of 0.1 mol/l. After that the energy was minimized using the Steepest Decent and Conjugate Gradient algorithms until machine precision was reached.

All simulations were done with GROMACS 2023.1 and 2023.4 ^92^. Coulombic interactions were handled using the Particle Mesh Ewald (PME) method with PME order of 4 and a Fourier spacing of 0.16. Close Coulomb real space interactions were cut off at 1.2 nm and Lennard-Jones (LJ) interactions were cut off after 1.2 nm without potential shift. Long-range dispersion corrections for energy and pressure were applied to account for errors stemming from truncated LJ interactions. Periodic boundary conditions were used. The Verlet neighbor-searching list with cut off 1.2 nm was used and updated every 10 steps. The leap-frog algorithm was used to integrate Newton’s equations of motion. A time step of 2 fs was used as hydrogen bonds were constrained using the LINCS algorithm ^93^.

A total of four different systems were modelled and simulated: The OAS2 monomer and dimer were simulated to investigate their conformational dynamics and judge their stability. They were simulated in cubic boxes of lengths 10.6 nm (monomer) and 11.4 nm (dimer). They contained approximately 146000 (monomer) and 192000 atoms (dimer). To locate the RNA binding site on OAS2 and asses the dynamics of the RNA-protein complex, a 44bp dsRNA was simulated together with the OAS2 monomer and dimer. These systems were set up in dodecahedral boxes with the dimensions (14.1 x 14.1 x 9.9) nm^3^ (monomer) and (16.5 x 16.5 x 11.7) nm^3^ (dimer) and contained around 260000 (monomer) and 421000 atoms (dimer).

### OAS2 monomer and dimer simulations

After minimization, the systems were equilibrated in the NVT ensemble for 4 ns at the physiological temperature of 310.15 K. All NVT equilibration simulations were done using the Berendsen thermostat ^94^ with a coupling constant of 0.1 ps. This was followed by 4 ns NPT equilibration, which employed the velocity rescaling algorithm ^80^ with a coupling constant of 0.1 ps and the Berendsen barostat using isotropic pressure coupling, a coupling constant of 2.0 ps, reference pressure of 1 bar and a compressibility of 4.5 *10^-5^ bar^-1^. Position restraints (1000 kJ/mol*nm^2^) were put on heavy protein atoms during NVT and NPT equilibration. After pre-equilibration the systems containing OAS2 monomer and dimer were simulated without restraints for 100 ns each in NPT ensemble, using the velocity rescaling thermostat with a coupling constant of 1.0 ps and the Parinello-Rahman barostat ^81^ with a coupling constant of 5.0 ps, reference pressure of 1 bar and a compressibility of 4.5 *10^-5^ bar^-1^.

### dsRNA binding site prediction simulations

For the predictions of the dsRNA binding site, the systems were simulated at a higher temperature to speed up the conformational sampling and to overcome possible energy barriers associated with binding and unbinding of the RNA. An initial temperature of 400 K was used. Once the RNA was bound, the systems were cooled to physiological temperature (see below).

For the OAS2 monomer and dimer, 10 simulation runs of 100 ns were initiated from the same equilibrated structures. The simulations were performed in the NVT ensemble at 400K using the velocity rescaling thermostat with a coupling constant of 0.1 ps. The starting velocities for each run were drawn randomly from Maxwell-Boltzmann. To prevent the RNA from unzipping at elevated temperatures, restraints were applied to the distances between the two terminal nucleobases at each end. Also, the position of the zinc ion within the binding site was restrained.

Potential binding structures were identified by analysing the number of close contacts between dsRNA and protein. For the dimer and monomer systems respectively, four potentially binding conformations were identified showing the maximum numbers of close contacts formed between domain I of the OAS2 monomer or dimer and the dsRNA. These structures were then taken as the starting points for additional equilibration at 310.15 K in NVT and NPT ensemble as described above. The pre-equilibrated dsRNA-OAS2 complexes were successively used for 100 ns production runs in NPT ensemble. The prediction of the key residues involved in the binding of dsRNA was based on these productions runs.

### Number of close contacts

The number of close contacts between dsRNA and OAS2 was calculated using GROMACS and the Contact Map Explorer [David W.H. Swenson and Sander Roet, https://github.com/dwhswenson/contact_map] Python package which builds on the MDTraj package ^95^. Close contacts were defined as instances where the distance between protein and RNA atoms was less than 0.35 nm. In line with the experimental findings of a minimal RNA length requirement for OAS2 activation, the analysis of close contacts was focused on domain I of OAS2.

### Contact frequencies

Contact Frequencies were calculated using the Contact Map Explorer [David W.H. Swenson and Sander Roet, https://github.com/dwhswenson/contact_map] Python package which builds on the MDTraj package ^95^ and additional python code. Close contacts were defined as protein and RNA atoms being closer than 0.35 nm. The frequencies were normalized in a way that a close contact of a protein residue with a dsRNA atom over the length of one whole 100 ns simulation resulted in a contact frequency of 1. Values above 1 either correspond to close contacts between a certain protein residue and multiple RNA atoms or one close contact that occurred for a length exceeding that of a single 100 ns simulation.

**The electrostatic surface potential** was calculated using the ChimeraX. The structure shown is based on one representative binding conformation extracted from one of the production runs. The structure shows the averaged backbone coordinates over a simulation time of 100 ns. The RMSD values were calculated as detailed below.

### RMSD

RMSD values were calculated using the Python library MDAnalysis ^96,97^ by aligning each frame of a 100 ns trajectory to the cryo-em structure and calculating the RMSD. For alignment and RMSD calculation only backbone atoms were considered. Figures showing the RMSD values by color were created using ChimeraX. They show structures averaged over 100 ns of simulation time. The corresponding RMSD values are based on alignments of the averaged structures obtained from MD simulations to the cryo-em structure. For the OAS2 monomer, half of the cryo-em dimer structure was used as a reference.

### Extended case report

The female patient was born at term as the first child of non-consanguineous healthy parents of Caucasian descent after an uneventful pregnancy. Her birth weight and length were normal and she developed normally. At the age of 4 years and 1 month, she presented with progressive fatigue since four weeks, epistaxis since two weeks, and hemochezia since one week. The pediatrician referred the child to the hospital because of a hemoglobin of 6.1 g/dl [11.1-14.3 g/dl]. On admission, acute-on-chronic renal failure was diagnosed with creatinine 4.4 mg/dl [0.2-0.6 mg/dl], urea 143 mg/dl [10-35 mg/dl], cystatin C 3.6 mg/dl [0.64-1.03 mg/l], metabolic acidosis (pH 7.29, base excess -9.8 mmol, HCO_3_ 17 mmol/l), and hyperkalemia 6.6 mmol/l [3.4-4.7 mmol/l]. Urine analysis showed microhematuria and nephrotic-range proteinuria (urine-protein/creatinine ratio 5590 mg/g). Infections with Hantavirus, CMV, EBV, HBV, HAV, HCV, HIV, streptococci, enterohemorrhagic E. coli, Campylobacter, Salmonella, and Shigella were excluded. She tested positive for perinuclear anti-neutrophil cytoplasmic antibodies (p-ANCA; 1:160 [<1:80]), antibodies to myeloperoxidase (MPO; 46 U/ml [<20 U/ml, and antinuclear antibodies (ANA 1:320 to 1:2000 [<1:80]]) (**Fig. 6A, SI Fig. 9A**). Renal ultrasound showed mildly hyperechogenic kidneys. Renal biopsy showed severe glomerulosclerosis (10 out of 18) with fresh and older areas of a proliferative extracapillary glomerulonephritis with two cellular and four fibrocellular crescents in eight non-scarred glomeruli and segmental necrotizing vasculitis of an interlobular artery, consistent with ANCA-associated vasculitis. There was also evidence of severe tubular epithelial damage with tubular atrophy and interstitial fibrosis, and chronic interstitial inflammation (**Fig 6B, Suppl Figure A, B**). Based on these findings, a diagnosis of MPO-ANCA-associated vasculitis with end-stage renal disease was made. Vasculitis was treated with i.v. methylprednisolone pulses (500 mg/m^2^ body surface area/day i.v. for 5 days), followed by prednisolone tapering (2.2-0.2 mg/kg body weight for 8 months), rituximab (2 doses á 375 mg/m^2^ body surface area) and azathioprine (2.8-1.5 mg/kg body weight/day for 3 months, stopped due to leucopenia). End-stage renal disease was treated by intermittent hemodialysis initially and switched to peritoneal dialysis. She developed severe hypertension and was treated with metoprolol, amlodipine, ramipril, and dihydralazine. The patient was also given vitamin D, calcitriol, calcium carbonate, azathioprine, potassium- and phosphate-reduced diet, erythropoietin, iron, and one blood transfusion. Hypertrophic osteoarthropathy was diagnosed 8 months after disease manifestation (**Suppl Figure C**). Because of increasing pANCA and MPO antibodies 11 months after disease manifestation, prednisolone was started again (0.6-0.1 mg/kg body weight/day). Growth retardation was attributed to an eating disorder, long-term corticosteroid therapy, and end-stage renal disease.

At the age of 5 years and 2 months, the patient was admitted to the hospital with fever and suspected bronchopneunomia. A chest x-ray showed bilateral infiltrations in the central lung areas extending to the lower fields. Tests for mycoplasma and SARS-Cov2 were negative. She was treated with ampicillin/sulbactam and discharged after five days. 10 days later, the patient presented with chest pain and dyspnea due to acute coronary syndrome with ST elevation myocardial infarction. Cardiac catheterization revealed non-perfusion of the anterior ramus interventricularis of the left coronary artery (**Suppl Figure D**). The patient was started on dual platelet aggregation inhibition and lysis therapy. Over the next two weeks, she experienced episodes of tachycardia arrythmia and ST elevations, a transfusion-dependent lower gastrointestinal bleeding, and hypertensive crises due to massive intracerebral hemorrhage with high intracranial pressure requiring craniectomy and deep sedation. Brain magnetic resonance imaging showed multiple bilateral ischemic areas in the occipital, left thalamus, and right temporal regions. A progressive increase in gamma-glutamyltransferase to 5900 U/l [<25 U/l] suggested primary sclerosing cholangitis as part of the vasculitis. After extubation, the patient developed hemiparesis of the right lower limb.

During acute coronary syndrome, the patient was found to have elevated inflammatory markers (**Fig 6A, SI Fig. 9A**). Despite negative investigations for infections including Influenza A/B, RSV, SARS-Cov2, Rotavirus, Adenovirus, Norovirus, HSV, VZV, and bacterial and fungal infections in tracheal secretions, blood, cerebrospinal fluid, and stool, she was treated with antibiotic therapy with piperacillin/tazobactam, tobramycin, and clindamycin. She was started on immunosuppressive therapy with mycophenolate mofetil (760 mg/m^2^ body surface area/day), one additional methylprednisolone pulse, and the prednisolone dose was again increased (up to 2 mg/kg body weight/day). She was also given immunomodulatory therapy with daily anakinra applications (for 3 weeks), one i.v. immunoglobulin administration, and infliximab (every four weeks) due to persistent inflammatory signs with elevated levels of C-reactive protein (CRP) up to 28.81 mg/dl [<0.5 mg/dl] and soluble IL-2 receptor (sIL-2R) up to 5808 U/ml [158-613 U/ml]).

At the age of 5 years and 7 months, she was readmitted for percutaneous endoscopic gastrostomy because of deterioration in her general condition and eating disorder with associated trichophagia and trichobezoar. During this hospitalization, the child developed epileptic seizures which were treated with midazolam and levetiracetam. She also developed signs of pulmonary edema with tachydyspnea and oxygen requirements, accompanied by elevated inflammatory markers (CRP 4.3 mg/dl [<0.5 mg/dl]; sIL-2R >6000 U/l [158-613 U/ml]). She was treated with methylprednisolone pulse therapy, midazolam and levetiracetam. Throat swabs were negative for Influenza/B virus, RSV, and SARS-Cov2. Due to progressive inflammatory signs, she was started on methylprednisolone pulses and prednisolone escalation, and switched from infliximab to anakinra (up to 5 mg/kg). Further investigations for autoinflammation revealed normal adenosine deaminase 2 activity and elevated interferon (IFN) signatures (IFN score 88.96 and 28.74 [<12.49]).

At the age of 6 years and 1 month, the child had an acute episode of chest pain, fatigue, and tachydyspnea. An ECG showed ST elevation and a cardiac echogram showed a massive decrease in cardiac contractility. Due to circulatory failure with hypotension, she was intubated and treated with catecholamines, hydrocortisone, and vasopressin. This was accompanied by a massive increase in interleuin-6 (IL-6 300,000 ng/l [<4 ng/l]), CRP 2.3 mg/dl [<0.5 mg/dl]) and leukopenia. Antibiotic therapy with meropenem and vancomycin was started. A cytokine storm due to bacterial sepsis was suspected as the cause of the circulatory failure, and she was also treated with methylprednisolone, anakinra, and immunoglobulins. However, myocardial function and circulation could not be stabilized due to refractory hypotension and inadequate organ perfusion, and the child died only few hours after admission to intensive care.

**Supplementary Figure.**
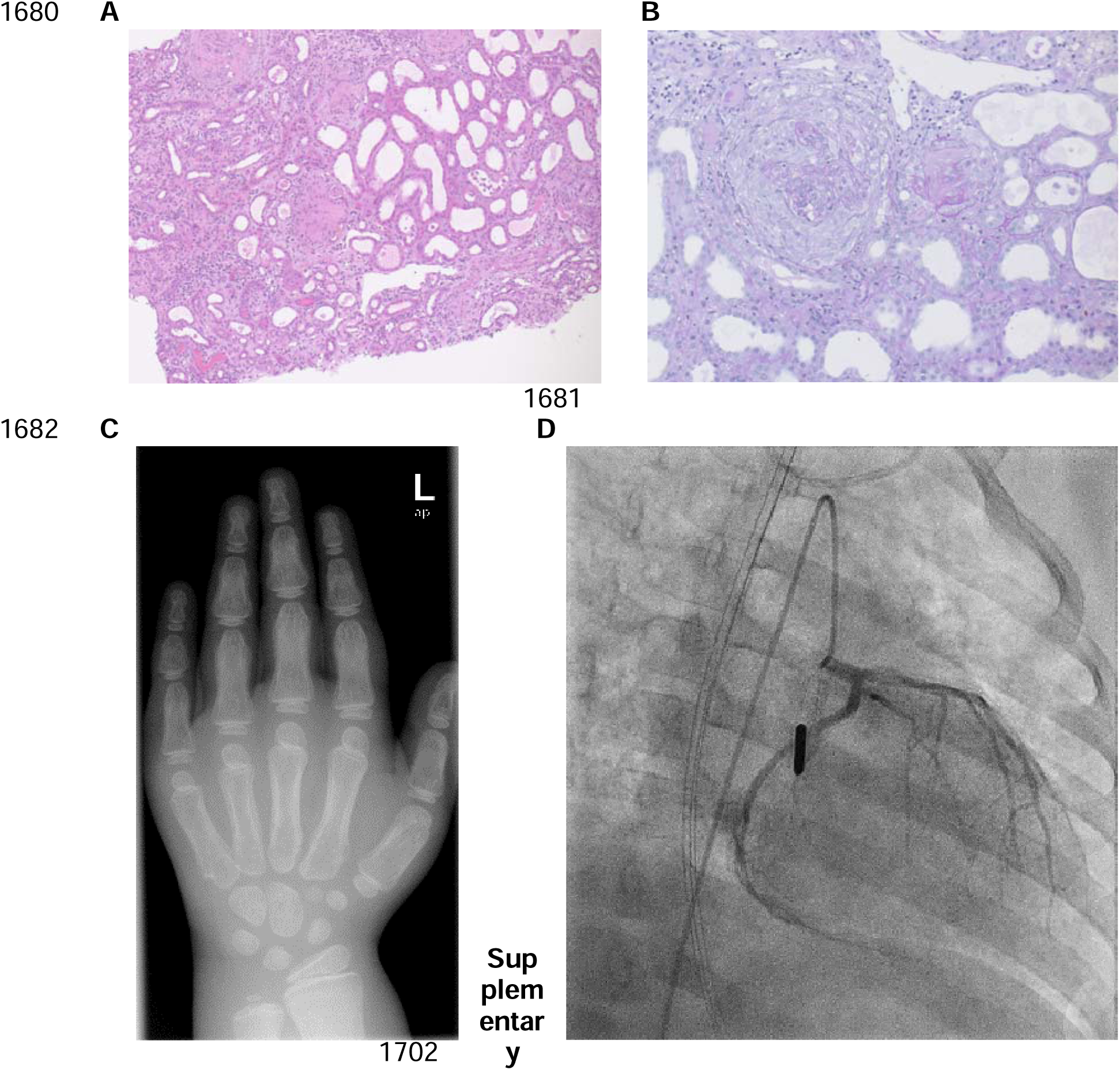
Renal histology, hand x-ray and coronary angiography. **A, B.** Histopathological findings of the renal biopsy showing areas of a proliferative extracapillary glomerulonephritis with cellular crescents, necrotizing vasculitis of an interlobular artery, along with tubular epithelial damage with tubular atrophy and interstitial fibrosis, as well as interstitial inflammation, consistent with ANCA-associated vasculitis. Hematoxylin and eosin stain, 10x magnification (A), periodic acid–Schiff stain, 20x magnification (B). **C.** X-ray of the left hand at the age of 5 years and 9 months showing osteopenic bone with double contours of the metacarpophalangeal and phalangeal periosteum, consistent with hypertrophic osteoarthropathy. **D.** Coronary angiography showing underperfusion of the peripheral segment of the left coronary artery, particularly the anterior interventricular ramus.

---

**Supplement Figure 1.**
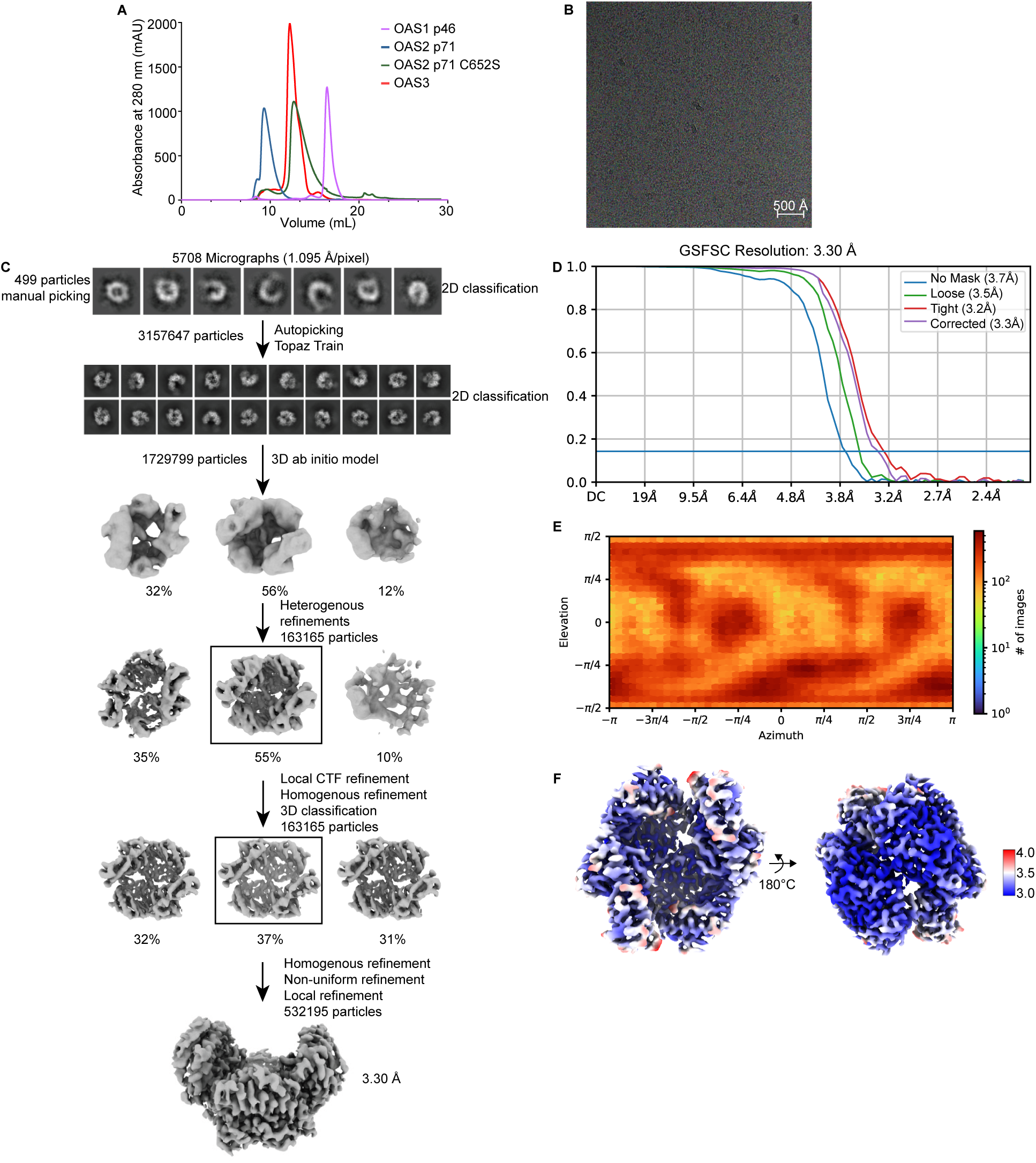
(A) Chromatograms of Superdex S200 10/300L of OAS1 p46, OAS2 p71, OAS2 p71 C652S and OAS3. (B) Representative micrograph of OAS2 apo data set. (C) Cryo-em data processing workflow of OAS2 apo data using cryoSPARC v4.5.3. (D) Map resolution estimated by Gold-standard Fourier shell correlation (GSFSC) with cutoff at 0.143. (E) Angular distribution dot plot made by CryoSparc Local refinement job. (F) Final reconstructed map colored by local resolution. Higher resolution is depicted in blue, lower resolution in red.

**Supplement Figure 2.**
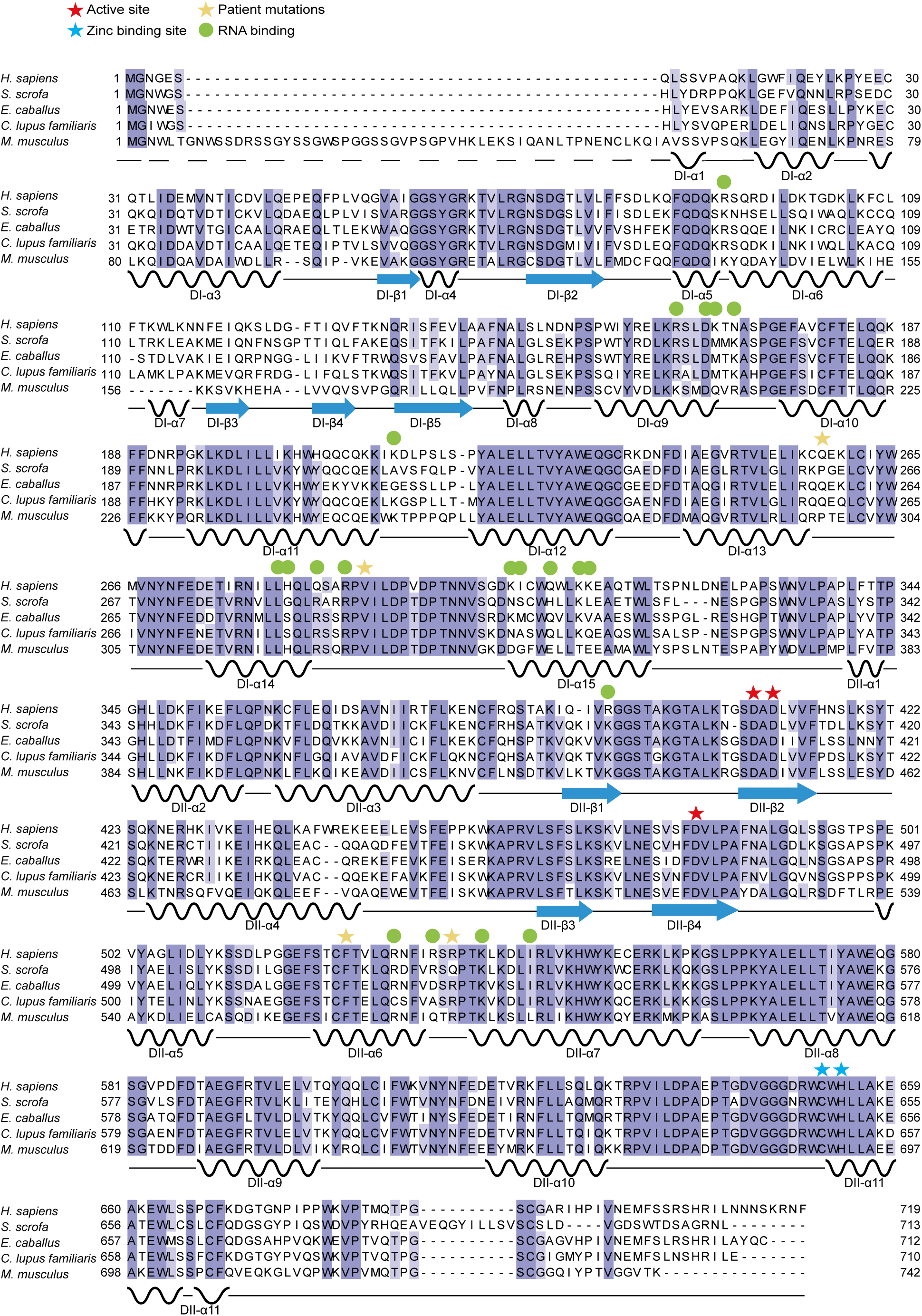
(A) Alignment of OAS2 from different species. Active site is depicted with red star, zinc binding site with blue star, RNA binding residues with green circle, patient mutation ^24^ with yellow star.

**Supplement Figure 3.**
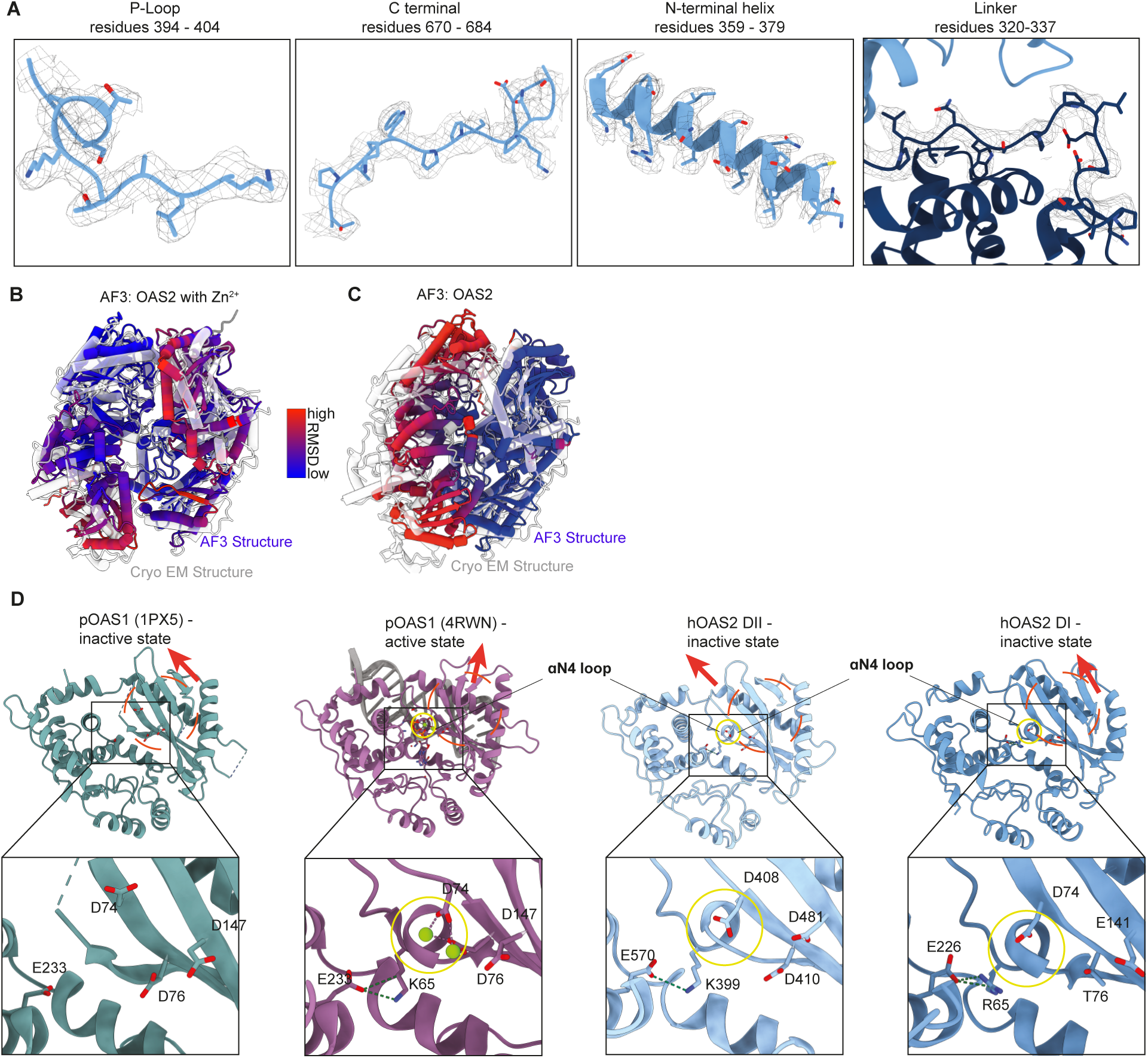
(A) Magnified density views of cryo-em map areas of P-loop, C-terminal end, N-terminal helix and Linker region. (B) Superposition of OAS2 cryo-em structure (grey) and Alphafold3 predicted dimer with Zinc ion colored by RMSD values with blue as low and red as high RMSD values. (C) Superposition of OAS2 cryo-em structure (grey) and Alphafold3 predicted dimer colored as (B). (D) Comparison of active site between pOAS1-dsRNA complex (PDB 4RWN, purple), OAS2 DII (light blue), OAS2 DI (dark blue) and pOAS1 apo structure (4RWQ). Yellow circles indicate location of P-loop. Red circles depict β-sheets, red arrows indicate orientation of β-sheets.

**Supplement Figure 4.**
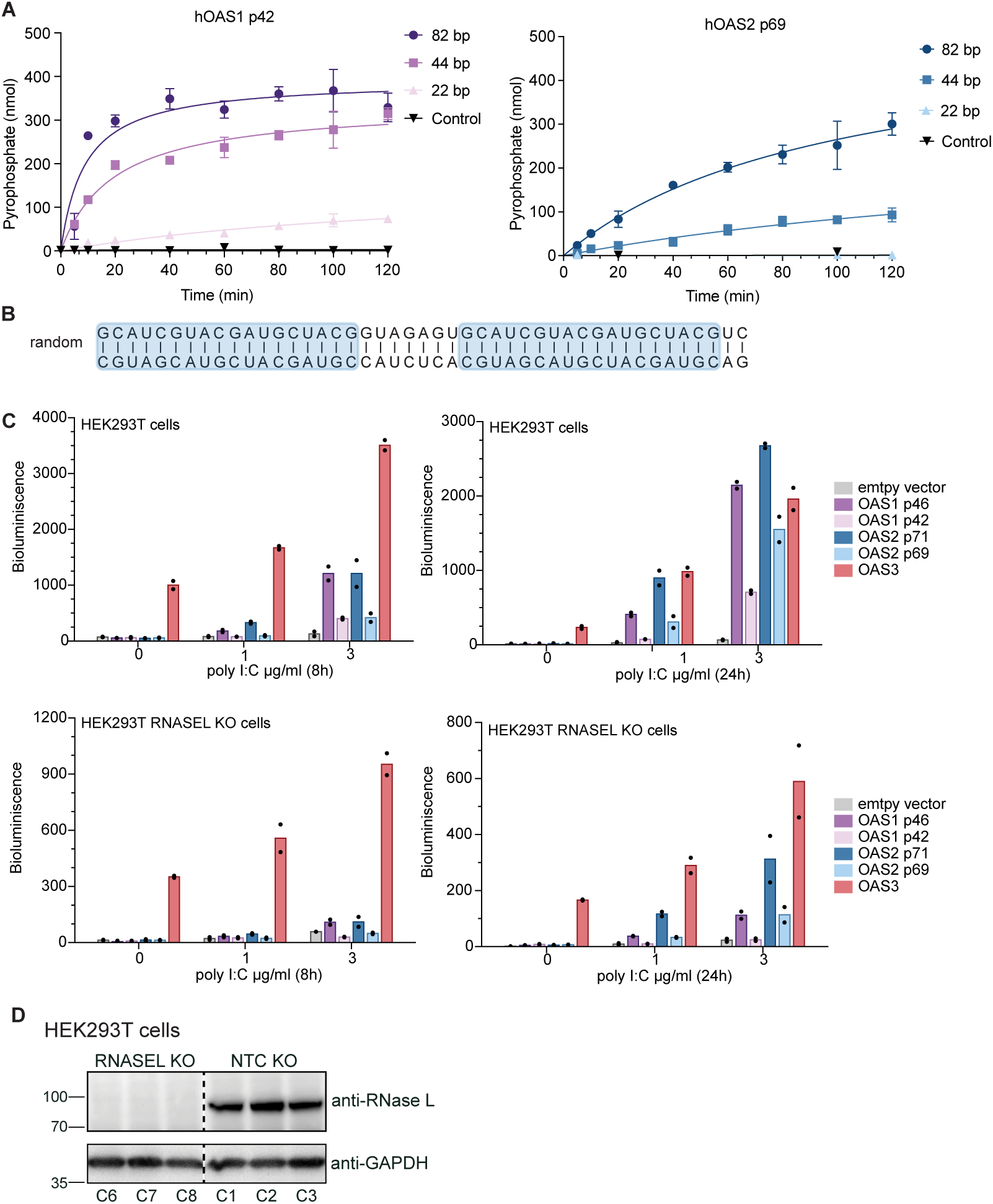
(A) In vitro chromogenic activity assay of 200 nM OAS1 p42 and OAS2 p69 with 22 bp, 44 bp, and 82 bp dsRNA (200 nM). dsRNA length is depicted in darker orange (long RNA) and light orange (short RNA) (mean ± SD of n = 3). (B) Sequence of random dsRNA shown in Fig. 2C. Repetitive sequence is depicted in blue. (C) Analysis of OAS protein activity in cells with 2′-5′OA biosensor for OAS1, OAS2 and OAS3 showing the highest activity for OAS3 even in the absence of poly I:C treatment. HEK293T or HEK293T RNASEL KO cells were transiently transfected with different OAS proteins for 48h followed by transfection of poly I:C. Bioluminescence was measured at 8h and 24h after poly I:C treatment. Assay is representative of at least three independent experiments. Bars represent means of technical replicates (dots). (D) Western blot validation of HEK293T RNASEL and NTC KO monoclonal cells used for 2’-5’OA biosensor assays.

**Supplement Figure 5.**
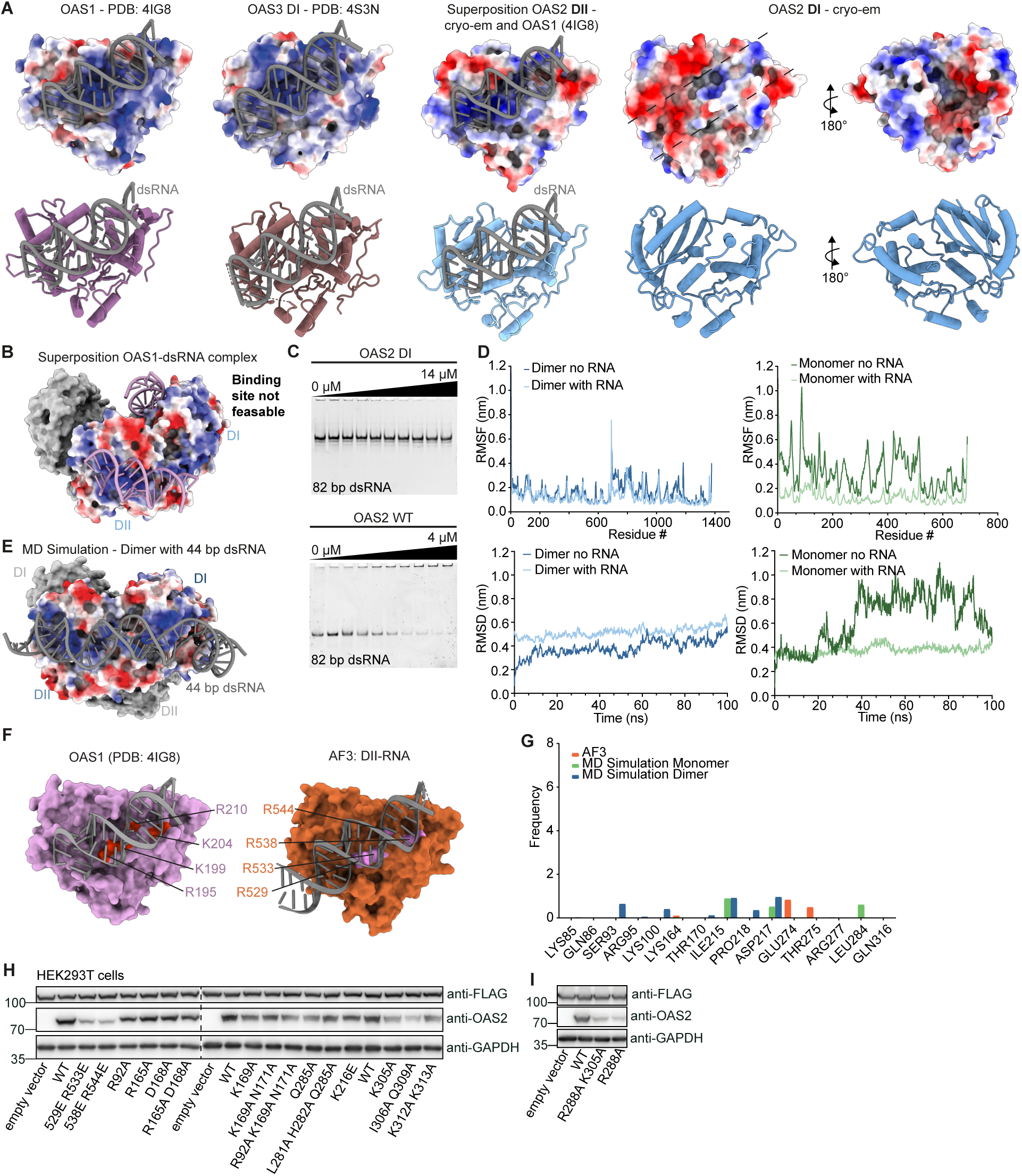
(A) Comparison of RNA binding sites of OAS1 (PDB 4IG8), OAS3 DI (PDB 4S3N), OAS2 DI and DII (Alphafold3 prediction) depicted as electron surface potential and ribbons. (B) Cryo-em structure of OAS2, one monomer depicted in grey, and one monomer depicted as electrostatic surface potential. Superposition of crystal structure of OAS1-dsRNA complex (PDB 4IG8, pink) with DI and DII shows predicted dsRNA binding sites on DI from top view (left) and DII from site view (right). (C) Electromobility shift assay of OAS2 DI and OAS2 WT with 82 bp dsRNA. Assays are representative of at least three independent experiments. (D) RMSF (top) and RMSD (bottom) plots of MD simulations. Simulation data of OAS2 dimer and monomer with RNA are depicted in blue and green, respectively. RMSD calculated after alignment of backbone atoms, reference: cryo-em structure. For RMSD and RMSF data are shown for monomer and dimer with and without RNA. (E) Electrostatic surface potential of MD simulation of OAS2 dimer in complex with dsRNA (grey). (F) Comparison of RNA binding interface of OAS1-RNA complex (4IG8) and AF3-DII. (G) Contact frequencies of AF3 predicted monomeric OAS2, MD simulated dimer and monomer with RNA for residues with low frequencies. (H) Western blot validation of OAS2 expression after transient transfection in HEK293T cells for analysis with 2’-5’OA biosensor. Anti-FLAG shows equal expression of FLAG-tagged reporter and anti-GAPDH was a loading control. (I) Western blot analysis of OAS2 wild-type and DI mutants with R288A mutations as in (H).

**Supplement Figure 6.**
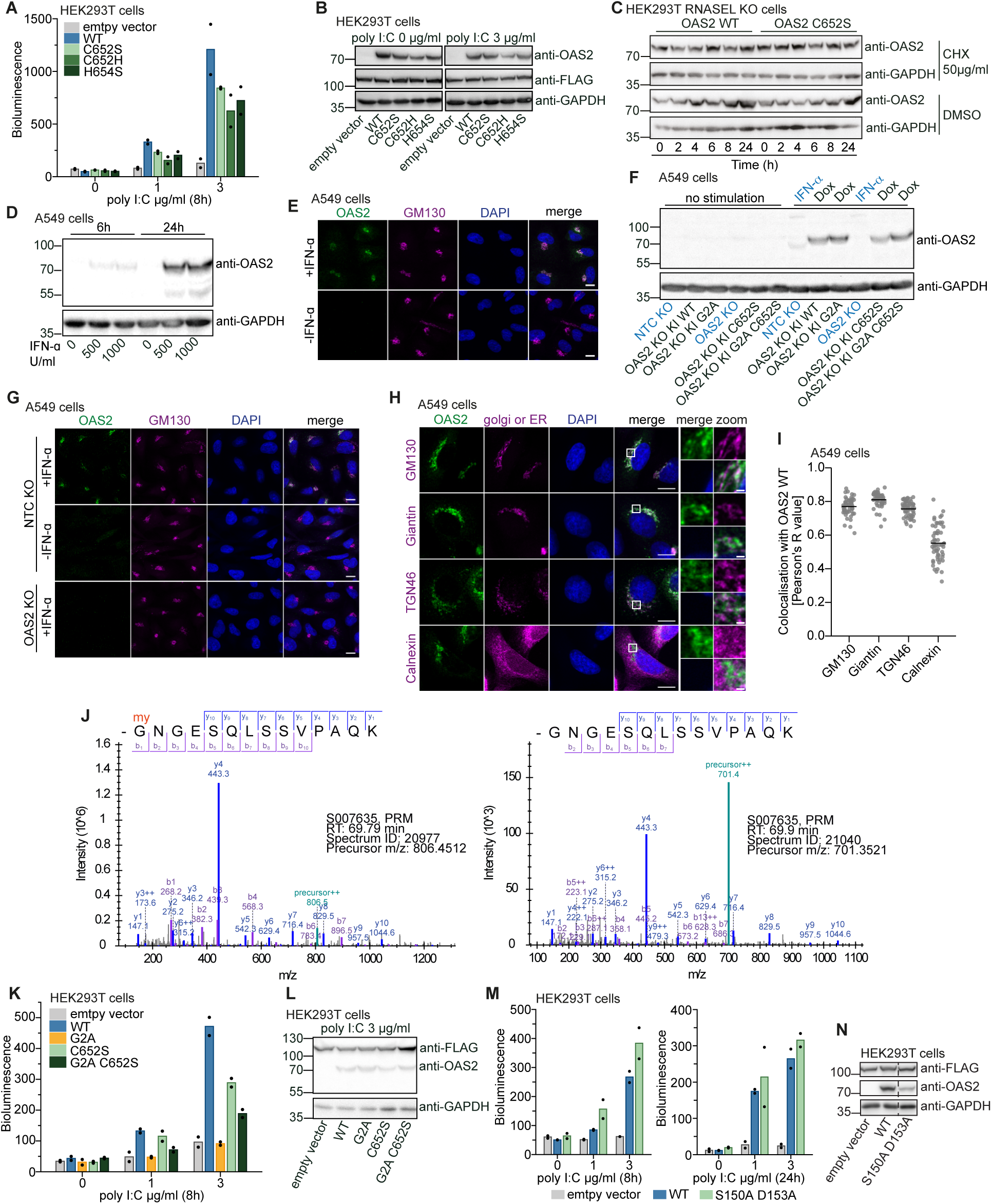
(A) Analysis of OAS2 activity in cells with 2’-5’OA biosensor for OAS2 wild-type and monomeric mutants showing higher activity of dimeric wild-type than the monomeric mutants. HEK293T were transiently transfected with different OAS2 constructs for 48h followed by treatment with poly I:C. Bioluminescence was measured at 8h after poly I:C transfection. (B) Western blot validation of OAS2 expression after transient transfection in HEK293T cells for activity measurements with 2’-5’OA biosensor. Anti-FLAG shows equal expression of FLAG-tagged reporter and anti-GAPDH was a loading control. (C) Western blot analysis of OAS2 WT and OAS2 C652S stability after cycloheximide (CHX) treatment. HEK293T cells were transiently transfected with OAS2 constructs for 24h followed by treatment with 50 µg/ml CHX or DMSO for indicated time points. (D) Western blot validation of endogenous OAS2 expression in A549 cells after stimulation with 500 U/ml or 1000 U/ml IFN-α for 6h and 24h. GAPDH is shown as a loading control. (E) Immunofluorescence confocal microscopy of OAS2 localization in A549 cells after stimulation with 500 U/ml IFN-α. Staining was performed for OAS2 (green), golgi marker GM130 (magenta) and DAPI (blue). Scale bars represent 10µm. (F) Western blot validation of OAS2 expression in A549 OAS2 KO cells reconstituted with doxycycline-inducible OAS2 constructs. Cells were stimulated with 500 U/ml IFN-α or 1 µg/ml doxycycline for 24h. GAPDH is shown as a loading control. (G) Immunofluorescence microscopy as in (E) in A549 OAS2 KO and NTC KO cells. (H) Immunofluorescence Airyscan microscopy of endogenous OAS2 localization in A549 cells after stimulation with 500 U/ml IFN-α for 24h. Staining was performed for OAS2 (green), DAPI (blue) and ER or golgi markers (magenta): GM130 (cis-golgi), giantin (medial-golgi), TNG46 (trans-golgi network) and calnexin (ER). Scale bars represent 10µm and 1µm for zoom in. (I) Quantification of colocalization of OAS2 and golgi or ER markers from (H) based on Pearson correlation. (J) Representative annotated ms/ms spectra for the myristoylated (left) and unmodified (right) GNGESQLSSVPAQK tryptic peptide of OAS2. Fragment ions of y-ion series (blue) and b-ion series (violet) are shown for each peptide. (K) Analysis of OAS2 activity in cells as in (A) for OAS2 wild-type and mutants with different localization showing that Golgi targeting is essential for the activity. (L) Western blot validation of OAS2 expression as in (B). (M) Analysis of OAS2 activity in cells as in (A) after 8h and 24h after poly I:C transfection for OAS2 wild-type and DI-DII interaction mutant S150A D153A showing higher activity when dimeric OAS2 wild-type conformation is destabilized in S150A D153A mutant. (N) Western blot validation of OAS2 expression as in (B). In (A), (K) and (M) assays are representative of at least three independent experiments. Bars represent means of technical replicates (dots).

**Supplement Figure 7.**
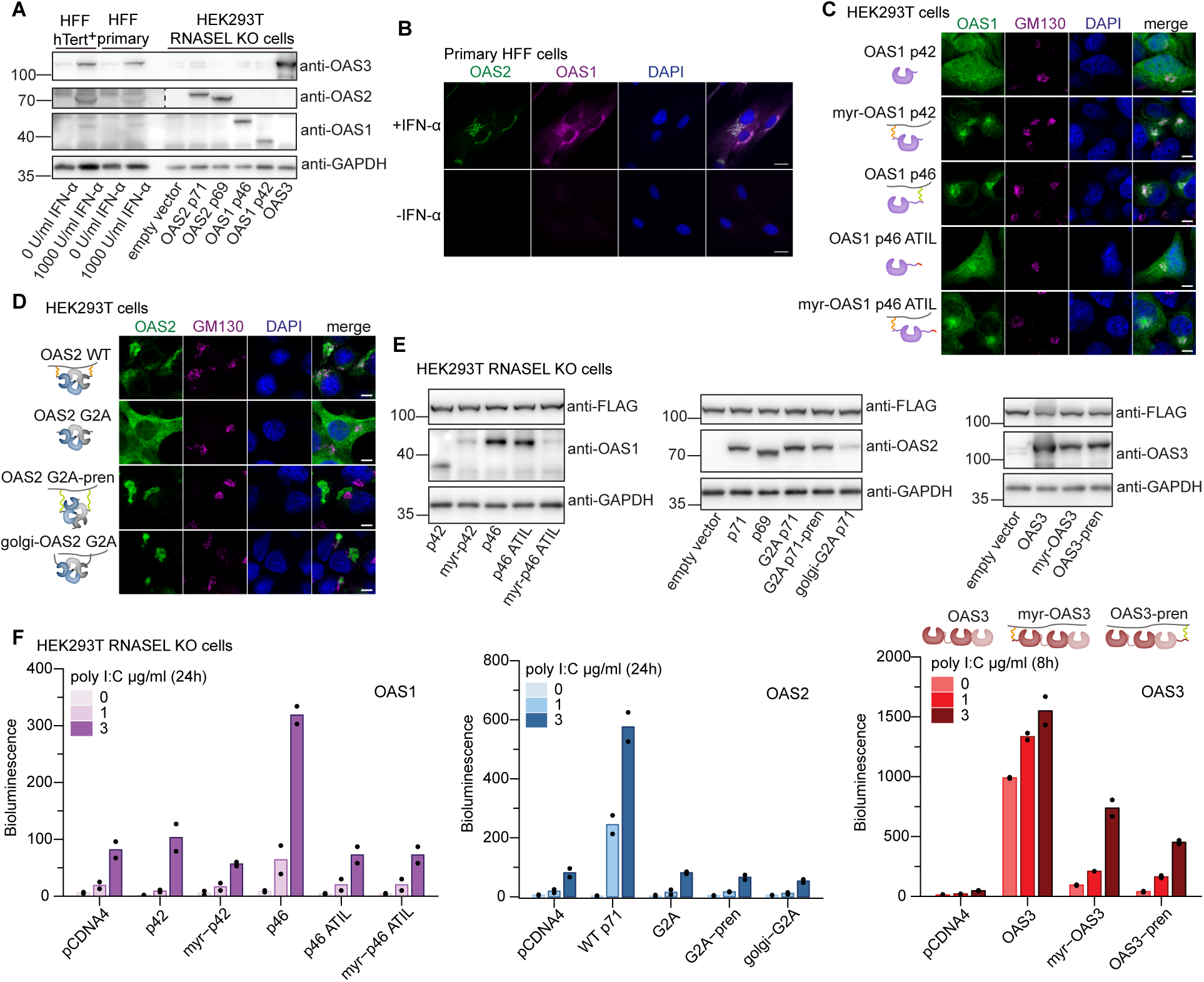
(A) Western blot analysis of OAS protein expression in BJ primary and hTERT^+^ HFF cells after stimulation with 1000U/ml IFN-α for 24h. Cell lysates from transient transfection of HEK293T RNASEL KO cells were loaded in parallel as a reference. GAPDH is shown as a loading control. (B) Immunofluorescence Airyscan microscopy of OAS2 and OAS1 localization in BJ primary HFF cells after stimulation with 1000U/ml IFN-α for 24h. Staining was performed for OAS2 (green), OAS1 (magenta) and DAPI (blue). Scale bars represent 20 µm. (C) Immunofluorescence Airyscan microscopy of OAS1 isoforms and chimeras in transiently transfected HEK293T cells. Staining was performed for OAS1 (green), golgi marker GM130 (magenta) and DAPI (blue). Scale bars represent 10 µm. (D) As in (C) but for OAS2. Golgi-OAS2 construct was designed by adding 10 amino acids to the OAS2 G2A N-terminus as shown in ^86^ to target OAS2 G2A to the Golgi. (E) Western blot validation for expression of OAS protein isoforms and chimeras in HEK293T RNASEL KO used for activity measurements in cells with 2’-5’OA biosensor in (F). Anti-FLAG shows equal expression of FLAG-tagged reporter and anti-GAPDH was a loading control. (F) Analysis of OAS2 activity in cells with 2’-5’OA biosensor for OAS protein isoforms and their chimeras with prenylation or myristoylation. Assay shows that OAS2 and OAS1 p46 are active only when myrisotylated and prenylated, respectively, whereas membrane targeting reduces the activity of OAS3. HEK293T were transiently transfected with different OAS2 constructs for 48h followed by transfection of poly I:C. Bioluminescence was measured at 24h (OAS1, OAS2) or 8h (OAS3) after poly I:C transfection. Assay is representative of three independent experiments. Bars represent means of technical replicates (dots). “myr” indicates N-terminal fusion with myristoylation motif from OAS2, while “pren” indicates C-terminal fusion with prenylation motif from OAS1 p46.

**Supplement Figure 8.**
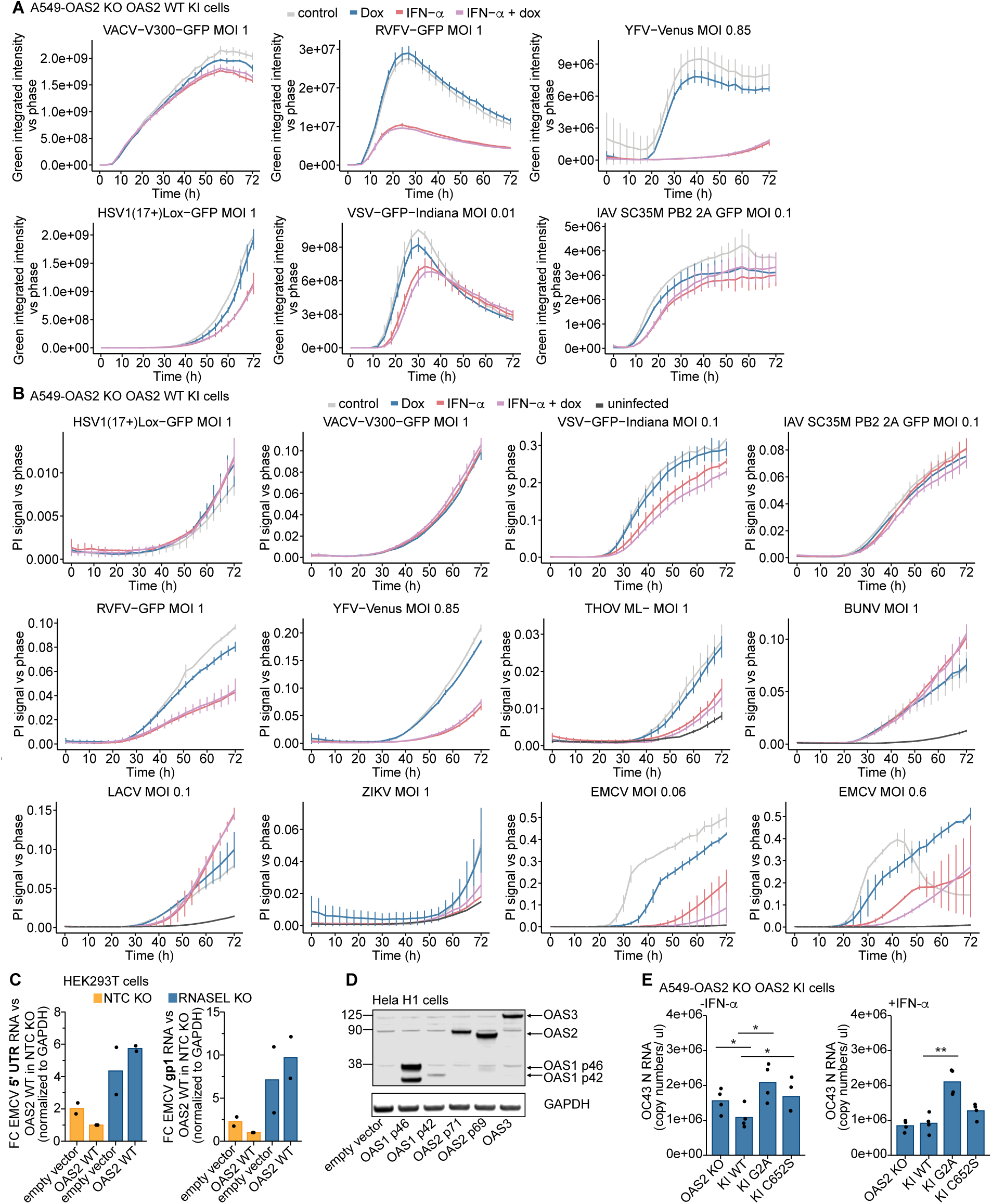
(A) Virus replication in A549 OAS2 KO cells reconstituted with doxycycline-inducible OAS2 WT after infection with different GFP-reporter viruses as described in Fig 5A. Virus replication was quantified as GFP integrated intensity versus cell confluency. (B) Cell death analysis as in (A). Cell death was quantified as area of propidium iodide signal versus cell confluency. (C) RT-qPCR analysis of intracellular EMCV RNA levels for 5’ UTR and gp1 in HEK293T monoclonal RNASEL KO and NTC KO cells. Cells were transfected with OAS2 WT or empty vector control for 24h followed by EMCV infection for 20h with MOI of 1. Bars represent means of 2 independent biological replicates (dots). (D) Western blot validation of OAS protein expression in Hela H1 cells after transient transfection. (E) RT-qPCR analysis of OC43 RNA levels in culture supernatants in A549 OAS2 KO cells reconstituted with doxycycline inducible OAS2 constructs. Cells were treated with 1µg/ml doxycycline with (right) or without 500U/ml IFN-β (left) for 24h followed by infection with OC43 for 72h. Bars represent means of four independent biological replicates (dots). Statistical significance was calculated using paired T-test. * p<0.05, ** p<0.001. In (A) and (B) data are plotted with error bars representing the SD of the mean from two technical replicates.

**Supplement Figure 9.**
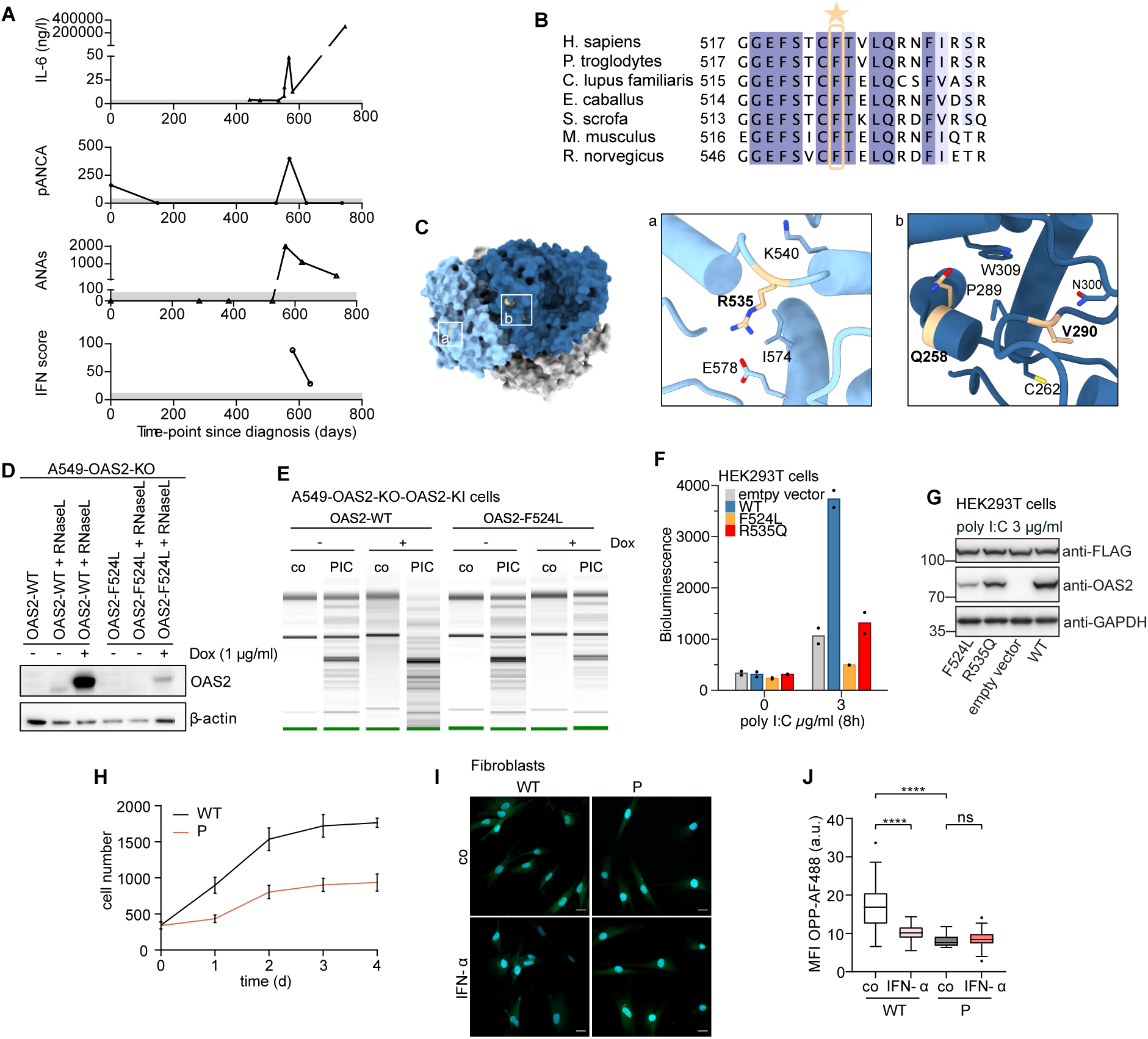
(A) Patient’s serum levels of interleukin-6 (IL-6), perinuclear anti-neutrophil cytoplasmic antibodies (pANCA), and antinuclear antibodies (ANA). The IFN score was calculated as previously described ^87^. Grey bars indicate normal ranges. (B) Multiple sequence alignment across different species indicating high conservation of the F524 of the human OAS2 protein. (C) Surface view of OAS2 depicted in two protomers, blue and grey. DI and DII are indicated in darker and lighter color, respectively. Patient mutations are indicated in yellow ^24^. Close up from (II) show interaction of residues Q258, V290 and R535. (D) Western blot validation of OAS2 expression in A549 OAS2 KO cells reconstituted with doxycycline-inducible OAS2 constructs. Cells were stimulated with 1µg/ml doxycycline for 24h and transfected with V6 RNase L WT 2’-5’OA biosensor. GAPDH is shown as a loading control. (E) RNA chip analysis of total RNA cleavage in A549 knockout cells reconstituted with either OAS2 WT or OAS2 F524L by doxycycline induction, untreated or treated with 2µg/ml poly(I:C) for 24 h. (F) Analysis of OAS2 activity in cells with 2’-5’OA biosensor for OAS2 wild-type and patient mutations F524L and R535Q showing loss of activity for the F524L mutant. HEK293T were transiently transfected with different OAS2 constructs for 48h followed by transfection of poly I:C. Bioluminescence was measured at 8h after poly I:C treatment. Assay is representative of at least three independent experiments. Bars represent means of technical replicates (dots). (G) Western blot validation of OAS2 protein expression from cellular activity assays with 2’-5’OA biosensor. Anti-FLAG shows equal expression of FLAG-tagged reporter. (H) Proliferation curve of wild-type (WT) and patient-derived fibroblasts (P) over 5 days. Data are mean ± SEM pooled from technical replicates. (I) Representative images of the OPP Alexa Fluo 488 protein synthesis analysis in untreated or IFN-□-stimulated fibroblasts from patient and healthy control, scale bar = 20 µm. (J) Quantification of mean fluorescence intensity (MFI) of Alexa Fluor 488-OPP in cells shown in (I). n = 26-55 cells. Box plots: center line, median; box, interquartile range; whiskers, 1.5x interquartile range. Two-way ANOVA with Sidak’s multiple comparisons test. **** p< 0.0001.

