## Supplemental Table 1 for "Structural Basis for OAS2 Regulation and its Antiviral Function"

### Table S1 Cryo-EM data collection, refinement and validation statistics

| | OAS2 apo (EMDB-51786) (PDB 9H1Z) |
|---|---|
| **Data collection and processing** | |
| Magnification | 130000 |
| Voltage (kV) | 300 |
| Electron exposure (e–/Å$^2$) | 50 |
| Defocus range (μm) | -1.4 – -2.9 |
| Pixel size (Å) | 1.059 |
| Symmetry imposed | C2 |
| Initial particle images (no.) | 2914103 |
| Final particle images (no.) | 532195 |
| Map resolution (Å) | 3.31 |
| 0.143 FSC threshold | |
| Map resolution range (Å) | 2.8 – 5 |
| **Refinement** | |
| Initial model used (PDB code) | AF2 AF-P29728-F1 |
| Model resolution (Å) | 3.48 |
| 0.5 FSC threshold | |
| Model resolution range (Å) | 80 – 3.2 |
| Map sharpening *B* factor (Å$^2$) | -84.47 |
| Model composition | |
| Non-hydrogen atoms | 10747 |
| Protein residues | 1323 |
| Ligands | 1 |
| *B* factors (Å$^2$) | |
| Protein | 77.42 |
| Ligand | 43.62 |
| R.m.s. deviations | |
| Bond lengths (Å) | 0.003 |
| Bond angles (°) | 0.562 |
| Validation | |
| MolProbity score | 1.59 |
| Clashscore | 6.36 |
| Poor rotamers (%) | 0 |
| Ramachandran plot | |
| Favored (%) | 95 |
| Allowed (%) | 5 |
| Disallowed (%) | 0 |
